# Massively parallel interrogation of human functional variants modulating cancer immunosurveillance

**DOI:** 10.1101/2024.04.25.591036

**Authors:** Ying Liu, Yongshuo Liu, Xuran Niu, Ang Chen, Yizhou Li, Ying Yu, Zhiheng Liu, Wensheng Wei

## Abstract

Anti-PD-1/PD-L1 immune checkpoint blockade (ICB) therapy has revolutionized clinical cancer treatment, while abnormal PD-L1 or HLA-I expression in patients can significantly impact the therapeutic efficacy. Somatic mutations in cancer cells that modulate these critical immune regulators are closely associated with tumor progression and ICB response. However, a systematic interpretation of cancer immune-related mutations is still lacking. Here, we harnessed the ABEmax system to establish a large-scale sgRNA library encompassing approximately 820,000 sgRNAs that target all feasible Serine/Threonine/Tyrosine residues across the human genome, which systematically unveiled thousands of novel mutations that decrease or augment PD-L1 or HLA-I expression. Notably, we revealed functional mutations that co-regulate PD-L1 and HLA-I expression, represented by the clinically relevant mutation SETD2_Y1666, and verified that it can benefit from immunotherapy *in vivo*. Our findings generate an unprecedented resource of functional residues regulating cancer immunosurveillance, meanwhile, offer valuable guidance for clinical diagnosis, ICB therapy, and the development of innovative drugs in cancer treatment.

## INTRODUCTION

In the course of cancer development and progression, tumors adopt diverse strategies to evade immunosurveillance and suppress antitumor immune responses, such as the activation of inhibitory checkpoints, dysfunction of antigen processing and presentation (APP), and editing of immunogenic neoantigens (Sharma et al., 2017; Spranger and Gajewski, 2018). Cancer immunotherapies, represented by ICB, have achieved remarkable efficacy in clinical trials for several malignancies. Among all immune checkpoints, the PD-1/PD-L1 pathway has stood out as an appealing target due to its therapeutic potential and relatively low immune toxicity. Multiple blockade antibodies have been approved for the treatment of various cancers, including melanoma, non-small cell lung cancer and renal cell carcinoma (Morad et al., 2021; Ribas and Wolchok, 2018). However, responses are limited to a small subset of patients, the underlined mechanisms remain to be fully elucidated (Kalbasi and Ribas, 2020; Ribas and Wolchok, 2018; Sharma et al., 2017). A series of tumor-intrinsic responsive hallmarks have been identified to impact the immunotherapy outcomes, especially involving the PD-L1 signaling pathway, major histocompatibility complex class I (MHC-I)-mediated APP, and interferon-γ (IFNγ) signaling in the tumor microenvironment (TME), whose regulation directly compromises antitumor activity and affects the efficacy of PD-1/PD-L1 blockade (Sun et al., 2018). Nonetheless, there is still a pressing need for a deeper understanding of the regulatory factors that influence both the response and resistance to ICB therapy.

Genetic screening has been extensively applied to target identification in cancer immunology. In recent years, several studies have employed CRISPR screens to uncover regulators of PD-L1 and MHC-I (HLA-I for human) in cancer cells, identifying numerous genes as functionally significant (Burr et al., 2019; Dersh et al., 2021; Gu et al., 2021; Mezzadra et al., 2017; Suresh et al., 2020). However, due to the limited resolution of canonical CRISPR/Cas9 screens, these approaches have primarily provided insights into the functional roles of regulators at the gene level. Somatic mutations in cancer cells, which can impact critical pathways related to immune regulation, are closely associated with clinical response to ICB treatment (Rooney et al., 2015; Shin et al., 2017; Zaretsky et al., 2016). Refer to the International Cancer Genome Consortium (ICGC) database, single-nucleotide variants are predominant and account for over 90% among varied types of somatic mutations, whose functional relevance remains poorly understood. With the development of base editing techniques, high-throughput functional screens based on base editing has revolutionized the canonical screening strategy, enabling to assess variant functions at the level of single amino acids or individual bases (Cuella-Martin et al., 2021; Hanna et al., 2021). A recent study used base editing screens to map mutations of key mediators of IFNγ pathway, providing an initial resource for understanding IFNγ signaling in cancer immune surveillance (Coelho et al., 2023). Nevertheless, a vast number of mutations with uncertain significance still require systematical investigation.

In clinical settings involving anti-PD-1/PD-L1 antibody treatments, the expression levels of PD-L1 or HLA-I on the cell surface of patients have been shown to have predictive value for ICB efficacy (Anderson et al., 2021; Havel et al., 2019; Kumagai et al., 2020; Montesion et al., 2021; Sun et al., 2018). Ongoing research has shown that post-translational modifications (PTMs) play pivotal roles in controlling PD-L1 expression and antigen presentation, through regulating the protein stability, translocation, and protein-protein interactions (Anderson et al., 2021; Cha et al., 2019). Among hundred types of PTMs, phosphorylation is the most common and extensively studied, primarily occurring on serine (S), threonine (T), and tyrosine (Y) residues in eukaryotes (Humphrey et al., 2015). Protein phosphorylation can broadly impact immune-related oncogenic or inflammatory signaling pathways, such as JAK/STAT, RAS, MAPK, and NF-κB pathways, thus affecting the anti-tumor immune response (Cha et al., 2019). However, despite the potential significance of phosphorylation sites, only a limited number of these sites have been thoroughly characterized.

In this study, we aimed to systematically identify critical sites involved in cancer immunosurveillance and the response to ICB therapy, focusing on potential phosphorylation sites on S/T/Y residues. Using an ABEmax-based sgRNA library coupled with the iBAR strategy (Zhu et al., 2019), we targeted all S/T/Y codons across the entire human genome. Through multiple high-throughput variant screens for regulators modulating PD-L1 and HLA-I expression, we identified thousands of novel residues within known regulatory genes and previously unknown genes, shedding light on their functional roles in the individual regulation and co-regulation for PD-L1 and HLA-I expression. Subsequently, we assessed the regulatory mechanisms of several candidate sites, including the clinically relevant mutation SETD2_Y1666, and proved their effects on enhancing ICB response in *in vivo* experiments. Our study provides an unprecedented resource of functional residues for understanding cancer immune response. Furthermore, the findings offer valuable insights for clinical diagnosis and the optimization of ICB treatment.

## RESULTS

### Genome-wide mapping of critical S/T/Y residues modulating PD-L1 expressions by ABE-based screening

The interaction between PD-L1 on tumor cells and PD-1 on T cells impedes activation, proliferation, and effector functions of antigen-specific CD8+ T cells, thus promoting cancer immune evasion (Sun et al., 2018). To systematically explore the functional residues modulating PD-L1, the core factor involved in immunotherapy, we leveraged ABEmax to generate site-directed mutagenesis for achieving large-scale screens. Our recent work has established an ABE-based sgRNA library targeting all feasible protein-coding regions containing S/T/Y residues within the editing window leading to missense mutations. This library encompasses a staggering 818,619 sgRNAs, which collectively target 277,051 S, 165,599 T, and 141,687 Y residues (a separate manuscript under review). The *de novo* synthesized S/T/Y library consists of two sub-libraries–one targeting the sense strand (465,554 sgRNAs) and the other one targeting the antisense strand (354,595 sgRNAs). Both sub-libraries were supplemented with the same negative controls targeting the *AAVS1* locus. To better handling such an extensive library effectively, the sgRNA library was constructed with three internal barcodes (iBARs) (hereinafter referred to as sgRNA^iBAR^ library), as previously described (Zhu et al., 2019). This system ensures a high-quality screening even at a high multiplicity of infection (MOI) while significantly reducing the number of cells required for the screening process.

PD-L1 expression can be driven by tumor-intrinsic mechanisms or induced by inflammatory cytokines, such as IFNγ, which is secreted by immune cells within the TME (Morad et al., 2021). To probe functional residues affecting cell surface PD-L1 expression in both constitutive and induced contexts, we performed screens using the S/T/Y sgRNA^iBAR^ library in a human melanoma cell line, A375, which was engineered to stably express ABEmax. This cell line exhibits low level of endogenous PD-L1 but shows substantial upregulation of PD-L1 upon exposure to IFNγ (Figure S1A). The two S/T/Y sub-libraries were separately transduced into A375-ABEmax cells at an MOI of 3. Subsequently, following ten days of sgRNA transduction, the library cells were subjected to both IFNγ-stimulated and non-stimulated conditions. Through two rounds of fluorescence-activated cell sorting (FACS) enrichment, we collected cell populations with either lower or higher level of surface PD-L1 expression in each condition (Figure 1A; Figure S1B-E). We also maintained a control group of library cells without FACS selection throughout the positive screening process. The library cells from the control group and FACS-selected experimental groups were subjected to next-generation sequencing (NGS), and the NGS data was subsequently analyzed using the MAGeCK-iBAR algorithm (Zhu et al., 2019). This analysis involved evaluating the change in sgRNA abundance and calculating the *p*-value for each sgRNA, considering the significance and consistency of three iBARs per sgRNA in each screen. The screen score was then generated as -log10 of the *p*-value after Benjamini-Hochberg (BH) adjustment (Figure 1A).

**Figure 1.**
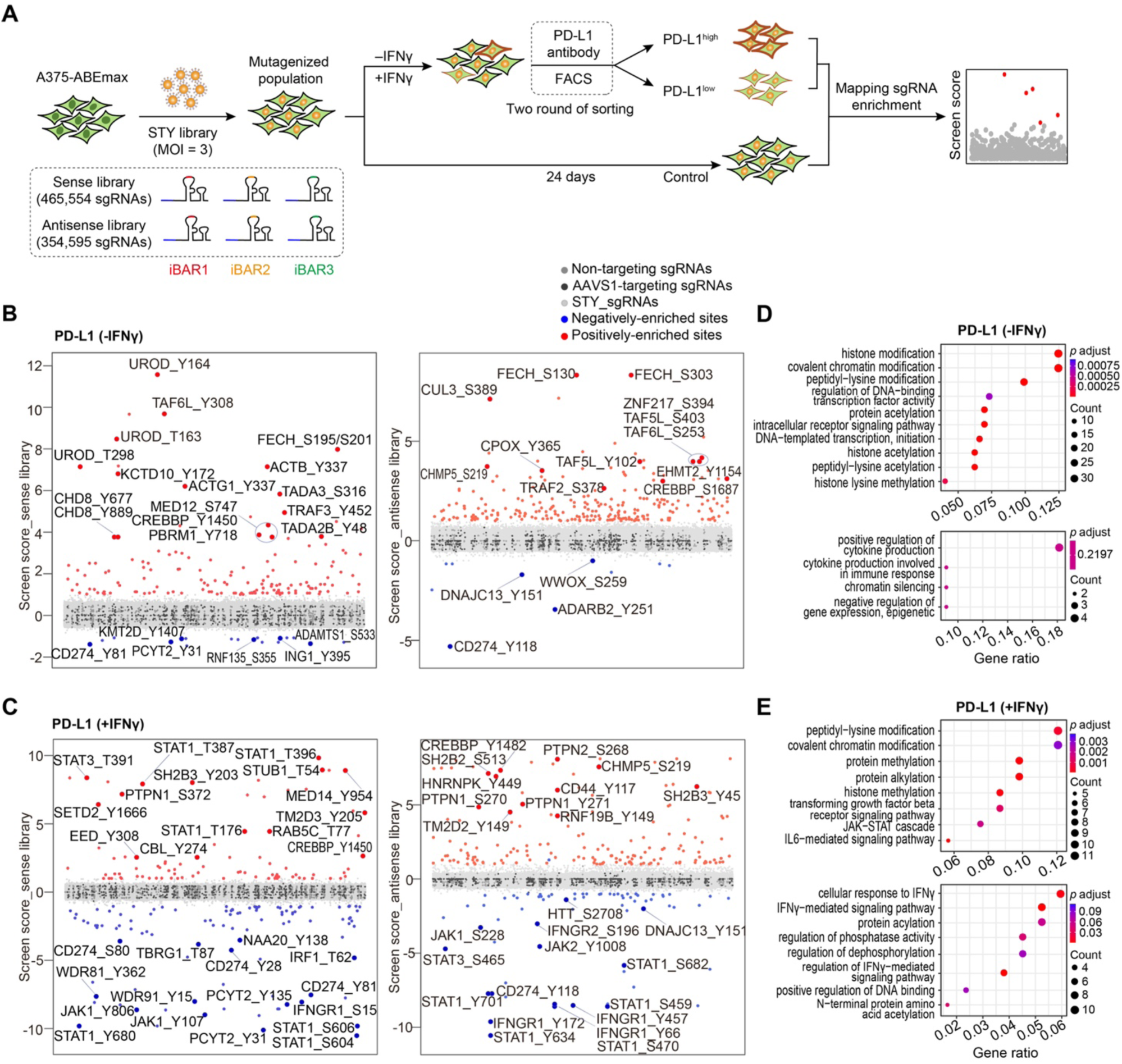
ABE-based screens identify functional S/T/Y residues modulating PD-L1 expression in genome-wide. (A) Schematic overview of the ABE screens for identifying S/T/Y residues that regulate PD-L1 expressions with and without IFNγ stimulation in A375 cells. (B-C) Significant S/T/Y residues enriched from the sense library (sense lib, left) and antisense library (antisense lib, right) that upregulate or downregulate PD-L1 expression in the absence of IFNγ (B) and upon IFNγ treatment (C). Positively or negatively enriched sites were selected by screen score > 1 or < -1. (D-E) Gene ontology (GO) enrichment analysis of related genes with identified mutations leading to PD-L1 upregulation (upper) and downregulation (lower) in the absence of IFNγ (D) and upon IFNγ treatment (E). See also Figure S1 and Table S1-4.

We selected sgRNAs with a screen score >1 for further investigations. In each screen, numerous novel sites were identified in both the high and low directions of regulating PD-L1 expression (Figure 1B-C; Table S1-4). To obtain a holistic understanding of the functional residues identified, we initially performed a gene ortholog (GO) analysis for all the related genes enriched in the screens, focusing on biological process. In the PD-L1 screen without IFNγ stimulation, the dominate terms in the PD-L1^high^ group were associated with histone modification, covalent chromatin modification, and the regulation of DNA-binding transcription factor activity. In contrast, the representative terms in PD-L1^low^ group included positive regulation of cytokine production and chromatin silencing (Figure 1D). In the PD-L1 screen with IFNγ exposure, the enriched terms were significantly correlated with interferon stimulation, encompassing processes such as the JAK-STAT cascade, transforming growth factor signaling pathway, cellular response to IFNγ, and the regulation of phosphatase activity. Moreover, some terms overlapped between the IFNγ-treated and IFNγ-absent conditions, particularly in PD-L1^high^ group, where terms such as peptidyl-lysine modification and covalent chromatin modification indicated the presence of conserved factors involved in tumor-intrinsic PD-L1 regulation, regardless of IFNγ treatment (Figure 1E).

### Massively parallel validation of regulatory variants affecting PD-L1 in A375 cells

For a deeper insight into the top-ranked hits from the screens, we integrated representative genes from both high and low directions in each screen. Subsequently, we built protein-protein interaction (PPI) networks using STRING followed by GO analyses. In IFNγ-absent PD-L1 screen, the network prominently showcased multiple genes enriched in processes related to histone modification, regulation of protein stability, chromatin remodeling, and heme biosynthesis process (Figure 2A). In the IFNγ-treated group, a large portion of genes were enriched in terms such as IFNγ-mediated signaling pathway, regulation of phosphorylation, and immune response (Figure 2B).

**Figure 2.**
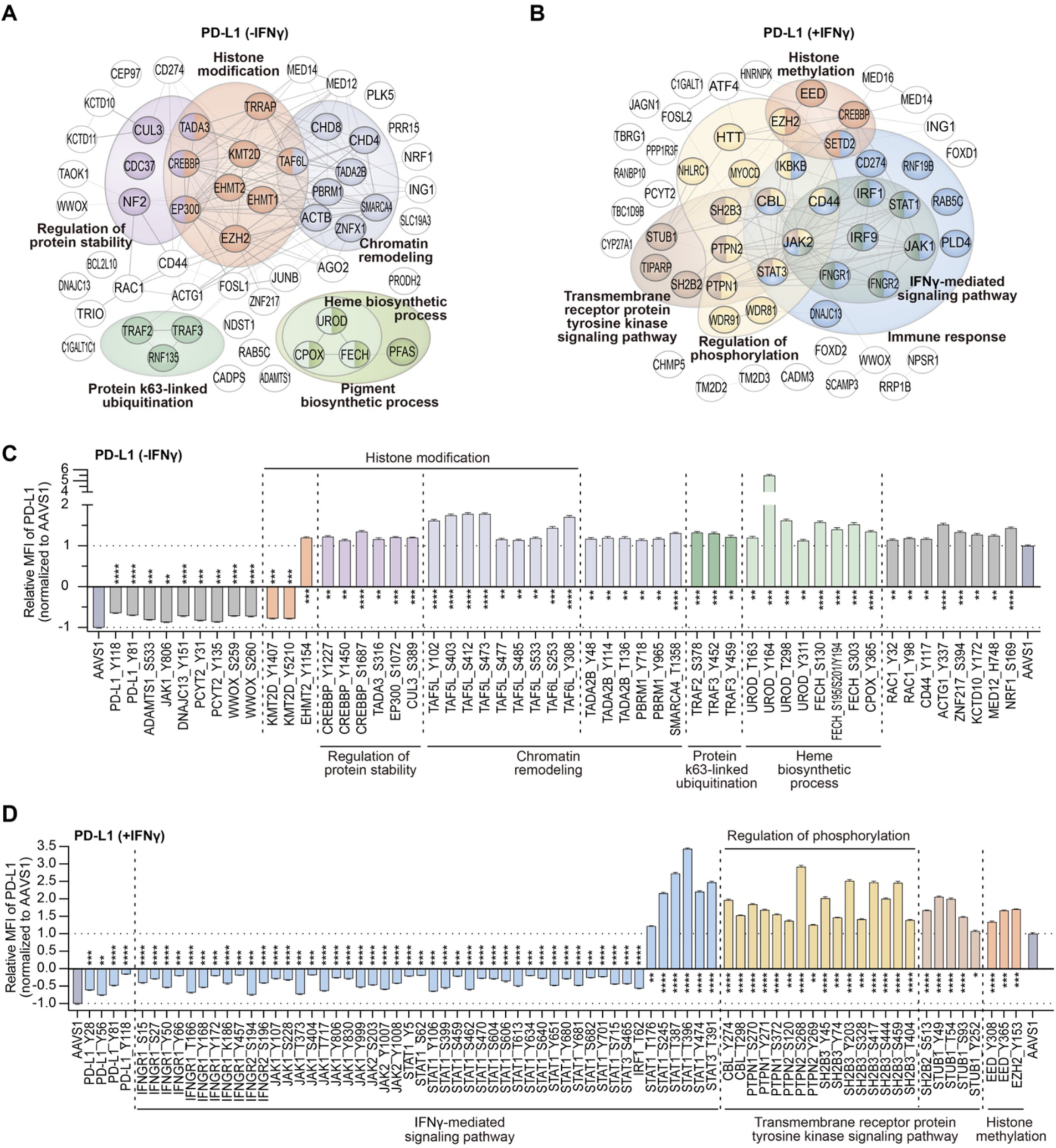
Validation of regulatory residues of PD-L1 enriched in various pathways. (A-B) STRING analysis of related genes with top-ranked mutations from IFNγ-absent PD-L1 screens (A) and IFNγ-treated PD-L1 screens (B). (C-D) Individual validations of negative and postive regulators of cell surface PD-L1 in A375 cells in the absence of IFNγ (C) and upon IFNγ treatment (D) by flow cytometry analysis. Cell surface PD-L1 was analysed following incubation without or with 100 ng/mL IFNγ for 48 h. The relative median fluorescence intensity (MFI) of surface PD-L1 for each mutant represents the ratio normlized to the MFI of *AAVS1*-targeting control cells. The data was presented as the mean ± SD (n=3). *P* values were calculated using two-tailed Student’s *t* test, \**P* < 0.05; ** *P* < 0.001; \*\*\**P* < 0.001; \*\*\*\**P* < 0.0001; NS, not significant. See also Figure S2 and Table S7.

To verify the regulatory roles of the identified variants, we selected candidate sites involved in different pathways and individually transduced each targeting sgRNA into A375-ABEmax cells via lentiviral infection. Subsequently, we conducted flow cytometry analysis to assess surface PD-L1 levels without or with IFNγ stimulation. Compared with the negative control sgRNA targeting the *AAVS1* locus, most of the sites showed significant regulation of PD-L1 expression.

In the absence of IFNγ, a standout performer was the UROD_Y164 site, alongside other confirmed residues within the UROD protein, including T163, T298 and Y311 (Figure 2C). UROD is involved in the heme synthesis pathway, whose disruption has been recognized to lead to an increase in PD-L1 expression (Suresh et al., 2020). Besides UROD, we also successfully verified the functionality of several mutations in FECH and CPOX, the other two core factors participating in heme synthesis but with no reported roles in regulating PD-L1 expression. Additionally, a series of novel sites enriched on genes associated with chromatin remolding, especially TAF5L and TAF6L, the integral components of the PCAF histone acetylase complex, were prominently ranked in the validation process (Figure 2C). Further analysis indicated that most of these variants showing a noteworthy phenotype influenced the expression of the target genes, ultimately resulting in an upregulation of overall and surface PD-L1 levels (Figure S2A). In the PD-L1^low^ group, due to its low baseline PD-L1 expression, a relatively smaller number of sites were identified and subjected to validation. Notably, PD-L1_Y118 and Y81 displayed the most significant impact, with Y118 being a previously recognized phosphorylation site. We also verified their association with PD-L1 expression for several additional sites, which are linked to genes known to be involved in immune response or ICB, such as WWOX_S259 and KMT2D_Y1407 (Chang et al., 2018; Wang et al., 2020) (Figure 2C).

Regarding the IFNγ-stimulated condition, several mutants reducing PD-L1 expression in IFNγ-absent condition were also validated under IFNγ treatment, including CD274_Y118, which exerted the strongest effect on downregulating surface PD-L1, consistent with the screening results (Figure 2D). Meanwhile, with IFNγ stimulation, more sites were identified and verified within these functional genes, such as *WWOX* and *PCYT2* (Figure S2B). A systematic analysis revealed that numerous mutations reduced the protein levels of their respective coding genes, as observed in the PD-L1^high^ group with *STUB1*, and in the PD-L1^low^ group with *WWOX*, *TBRG1*, and *IKBKB* (Figure S2C-D). Moreover, there were variants that did not significantly affect their protein expression, including HNRNPK_Y449, EED_Y308, and EED_Y365, suggesting that they may induce PD-L1 expression through other mechanisms (Figure S2C). Remarkably, a substantial number of sites were enriched on genes linked to the IFNγ-mediated signaling pathway and regulation of phosphorylation (Figure 2D; Figure S2B), we thus delved into investigating the regulatory mechanisms of these candidate sites.

### Systematic combing of functional residues within the IFNγ signaling pathway

We observed plenty of novel sites emerged on well-established genes linked to the IFNγ-mediated signaling pathway, including IFNγ receptors, Janus kinases, among others. Most of these mutations negatively regulated PD-L1 expression, affecting their respective coding genes, such as *IFNGR1*, *IFNGR2*, *JAK1*, *JAK2*, and *IRF1*. All investigated mutations on IFNGR1 and IFNGR2 were found to simultaneously decrease the membrane and overall PD-L1 protein levels (Figure 2D; Figure S3A). Notably, IFNGR1_Y457, a known phosphorylation site responsible for mediating the interaction between IFNGR1 and STAT1 proteins (Qing et al., 2005), was found to significantly downregulate PD-L1 expression. This suggests that the IFNGR1_Y457H mutation might affect its binding with STAT1, blocking the transmission of IFNγ signals and resulting in a substantial reduction in PD-L1 expression. Similarly, mutations in the downstream non-receptor tyrosine kinases JAK1 and JAK2 generally led to a decrease in the overall protein level of PD-L1 (Figure S3B-C). Meanwhile, two mutations on JAK2 consistently reduced both the mRNA and protein levels of JAK2, while most verified sites on JAK1 did not affect its own expression at both the mRNA and protein levels (Figure S3C-D). In addition, multiple sites on JAK1 and JAK2 were closely related to phosphorylation, as exemplified by four known phosphorylation sites and two predicted phosphorylation sites on JAK1, and two conserved phosphorylation sites, Y1007 and Y1008, on JAK2, which are critical for JAK2 function (Lucet et al., 2006) (Figure S3B).

Intriguingly, some genes related to the IFNγ signaling pathway contained residues with both negative and positive regulatory roles in PD-L1 expression. Notably, *STAT1* and *STAT3* were identified in this context (Figure 2D), which could not be detected in canonical screens at the gene level. STAT1, an important transcription factor connecting cytokine receptors with downstream target genes, is involved in the signaling of many cytokines, including IFNγ. In our screening, numerous functional S/T/Y sites were identified on the STAT1 protein, distributed across its four domains as well as the coiled-coil region (Figure 3A). Among them, five mutations were confirmed to upregulate PD-L1 expression, with two in the coiled-coil region and three in the DNA binding domain. The majority of mutations appeared to inhibit PD-L1 expression and were dispersed across functional regions, including the N-terminal domain, DNA-binding domain, SH2 domain, phosphorylated tail segment, and the transcriptional activation domain. One of the well-known sites was STAT1_Y701, located in the phosphorylated tail segment, where phosphorylation is required for the dimerization and nucleation of STAT1 (Quelle et al., 1995). Besides Y701, we also identified another confirmed phosphorylation site, STAT1_Y106, and 9 predicted phosphorylation sites that resulted in decreased PD-L1 expression following mutation, among which 7 sites were in the SH2 domain, indicating a close relationship between the SH2 domain and phosphorylation-mediated signaling transmission.

**Figure 3.**
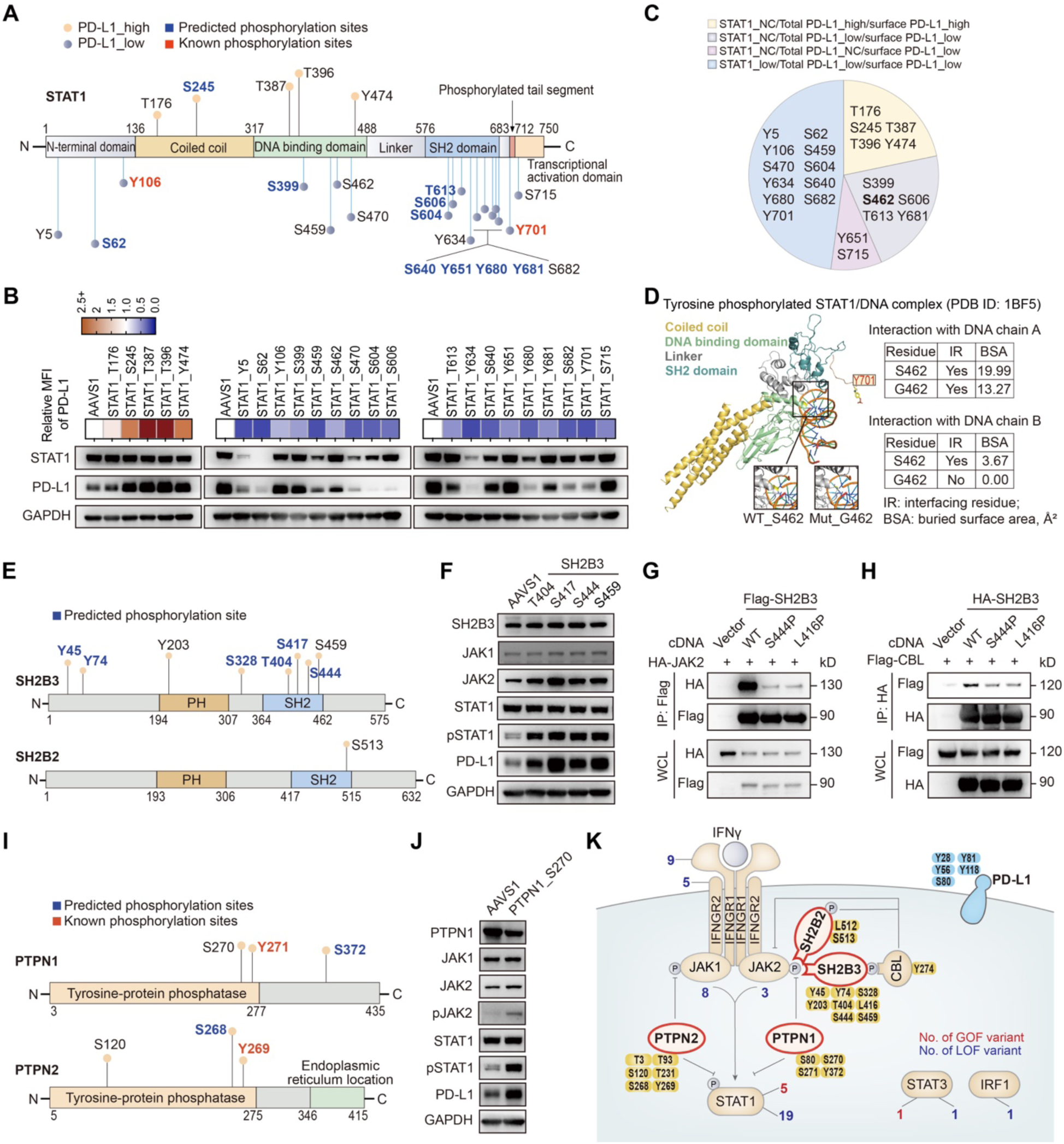
Novel residues on canonical and non-canonical regulatory proteins involved in IFNγ signal transduction affect the expression of PD-L1. (A) Distribution of identified S/T/Y residues on STAT1 protein. The regulatory residues are marked in two directions on the protein structure, with mutations upside indicating negative regulators and mutations downside indicating positive regulators. The relative length of each vertical line reflects the regulatory effect of the indicated residue according to the results of flow cytometric analysis from Figure 2D. (B) Protein expression levels of STAT1 and PD-L1 in the indicated A375 mutant cells treated with IFNγ. The upper heatmap shows the relative surface PD-L1 level of A375 cells with each mutation according to the results of flow cytometric analysis from Figure 2D. The lower IB analysis shows the total protein level of STAT1 and PD-L1 for each corresponding mutant. (C) Pie chart of STAT1 residues that are classfied based on the differential regulation of STAT1, total PD-L1 and surface PD-L1 expression. (D) Schematic of the molecular structure and intramolecular interactions around STAT1_S462 residue within tyrosine phosphorylated STAT1 and DNA complex (PDB: 1BF5). The WT S462 or the mutated G462 residue is labeled in yellow (left). The table shows the interaction between STAT1_S462/G462 with DNA chain A/B, which is indicated by the parameters of interfacing residue (IR) and buried surface area (BSA) (right). (E) Distribution of identified S/T/Y residues on SH2B2 and SH2B3 proteins. The regulatory residues are marked above each protein structure, which indicate negative regulators. The relative length of each vertical line reflects the regulatory effect of the indicated residue according to the results of flow cytometric analysis from Figure 2D. (F) IB analysis of typical JAK/STAT signaling components, SH2B3, and PD-L1 in A375 cells infected with respective sgRNA targeting *AAVS1* and each mutation. (G) IB analysis of anti-Flag immunoprecipitates (IPs) and whole-cell lysates (WCLs) of 293T cells co-transfected with the indicated plasmids expressing HA-tagged JAK2 and Flag-tagged SH2B3 WT or variants. (H) IB analysis of anti-HA IPs and WCLs of 293T cells co-transfected with the indicated plasmids expressing Flag-tagged CBL and HA-tagged SH2B3 WT or variants. (I) Distribution of identified S/T/Y residues on PTPN1 and PTPN2 proteins. The way of labeling each residue is the same as Figure 3A. (J) IB analysis of typical JAK/STAT signaling components, PTPN1, and PD-L1 in A375 cells infected with respective sgRNA targeting *AAVS1* and PTPN1_S270. (K) Schematic diagram of SH2-B family proteins and PTP family proteins regulating IFNγ-induced JAK/STAT signaling pathway. The information of identified S/T/Y residues were labeled on related proteins. The number of negative regulators are labeled in red and positive regulators are labled in blue. All cell samples were treated with 100 ng/mL IFNγ for 48 h. See also Figure S3 and Table S7-8.

We further performed immunoblot (IB) verification for all the selected sites within STAT1. Five PD-L1^high^ variants consistently upregulated PD-L1 expression in both total and membrane protein levels, while leaving the STAT1 protein level unchanged (Figure 3B). We hypothesized that these variants represent gain-of-function (GOF) mutations that promote the shuttle of STAT1 into the nucleus, facilitating its binding to DNA. Conversely, the majority of PD-L1^low^ mutations, distributed across various domains of STAT1, had an inhibitory effect on STAT1 expression, resulting in a decrease in both total and membrane protein levels of PD-L1. Interestingly, nearly half of these variants had no discernible impact on STAT1 expression. Some of them only affected the membrane PD-L1 levels, leaving the total PD-L1 level unchanged. This category includes mutations such as Y651 and S715. Others induced PD-L1 reduction in both total and membrane protein levels (Figure 3C). One notable example in this category is STAT1_S462G, a variant located in the DNA binding domain of STAT1, which was speculated to destroy the interaction between STAT1 and DNA. To verify this conjecture, we investigated the interaction between STAT1 and DNA before and after S462 mutation using the PDBePISA website (https://www.ebi.ac.uk/pdbe/pisa/). The analysis revealed that STAT1_S462 has interface contact with both strands of DNA, indicating that S462 is located at the interaction interface. However, S462G mutation completely abolished the ability of STAT1 to interact with one strand of DNA and decreased the contact area with the other DNA strand (Figure 3D). The analysis indicated that S462G mutation is likely to reduce the binding capacity of STAT1 to DNA, weakening its transcriptional effect and ultimately affecting the expression level of PD-L1.

### Critical S/T/Y residues within adaptor proteins and tyrosine phosphatases participate in the regulation of PD-L1 expression

In addition to the proteins directly involved in the IFNγ signaling pathway mentioned above, a series of mutations were associated with the regulation of phosphorylation. Among them, multiple residues on two types of proteins, which belong to the Src homology 2-B (SH2-B) protein family and the protein tyrosine phosphatase (PTP) protein family, were significantly enriched in the PD-L1^high^ screen. It’s worth noting that, for most of these proteins, their relevance in PD-L1 regulation especially at the residue level has not been intensively investigated in previous studies.

The SH2-B family, comprising SH2B1, SH2B2 (APS), and SH2B3 (Lnk) (Ahmed and Pillay, 2001), is a conserved family of adaptor proteins with similar structural characteristics. They possess a Pleckstrin homology domain (PH) that recognizes phosphatidylinositol lipids, enabling protein transfer to the cell membrane, as well as an SH2 domain that recognizes phosphorylated tyrosine residues (Figure 3E). In human cells, SH2-B proteins recognize and bind to phosphorylated Y813 of JAK2 via their SH2 domains (Bersenev et al., 2008; Kurzer et al., 2006), and an active region at the C-terminal of these proteins gets phosphorylated, interacting with the tyrosine kinase binding (TKB) domain of the intracellular E3 ubiquitin ligase CBL (c-cbl). This interaction recruits CBL to the vicinity of JAK2, leading to the degradation of JAK2 through ubiquitination modification, thereby negatively regulating the IFNγ signaling pathway (Hu and Hubbard, 2005).

We found that the residues with the most significant effects were clustered in the SH2 domain of these proteins, one for SH2B2 and four for SH2B3 (Figure 3E). For SH2B3, all four mutations did not alter the protein level of SH2B3 but significantly increased the total abundance of PD-L1 protein.

These mutations were found to activate the IFNγ-induced JAK-STAT signaling pathway, with JAK2 showing increased abundance and pSTAT1 levels significantly elevated (Figure 3F). The overall pattern of SH2B2_S513 closely resembled that of SH2B3 (Figure S3E), suggesting that both of these proteins influence JAK-STAT signaling by regulating the protein abundance of JAK2. Additionally, CBL_Y274 was identified and verified in the study, which located within the TKB domain and closely related to the recognition of SH2-B family (Hu and Hubbard, 2005). Its regulation on downstream JAK-STAT signaling was consistent with that of SH2B2 and SH2B3 (Figure S3F), further highlighting the critical role of the "JAK2-adaptor-CBL" loop in regulating IFNγ-mediated JAK/STAT signaling pathway and PD-L1 expression.

To further understand the regulatory mechanisms of these mutations, we focused on the SH2 domain and selected representative sites, namely SH2B3_S417, S444, and SH2B2_S513, for further investigation. Genomic sequencing indicated that targeting SH2B3_S417 generated L416P mutation, SH2B3_S444 generated the expected S444P mutation, and SH2B2_S513 targeting generated S513P and the bystander mutation L512P (Supplemental Information). Consequently, we overexpressed both the wild-type (WT) cDNAs and all the corresponding mutant sequences of these two genes to perform co-immunoprecipitation (Co-IP) experiments with JAK2 and CBL, respectively. Both the L416P and S444P mutations in SH2B3 simultaneously disrupted the interaction between SH2B3 and JAK2, as well as CBL, with a particularly notable impact on the interaction with JAK2 (Figure 3G-H). This severe destruction in the interactions with both JAK2 and CBL resulted in a weakened ubiquitination degradation of JAK2, leading to JAK2 upregulation and enhanced IFNγ signal, ultimately promoting PD-L1 expression. Intriguingly, distinct from the residues in SH2B3, none of the SH2B2_L512P, L513P single mutant, and L512P/S513P double mutant affected the interaction between SH2B2 and JAK2, but significantly reduced the interaction between SH2B2 and CBL (Figure S3G-H). We speculated that SH2B3_L416 and S444 are located close to the interaction center where the SH2 domain binds to JAK2_Y813 (Hu and Hubbard, 2005), while not for SH2B2_S512 or S513, thereby leading to a clear disruption in the interaction between SH2B3 and JAK2 after mutation. The analysis above suggests that mutations, especially those occurring within the SH2 domain of these two adaptor proteins, can dramatically affect the IFNγ signaling pathway, albeit through different regulatory patterns.

The screen also identified functional residues within PTPN1 and PTPN2, two members of the PTP family known to negatively regulate the cytokine signaling pathway through dephosphorylation of phosphorylated tyrosine residues on targeted proteins (Gu et al., 2003; Kleppe et al., 2011). Most of the identified residues were enriched within the phosphatase domain of each protein (Figure 3I). As such, we speculated that these mutations might affect their phosphatase activity, thereby enhancing the transmission of IFNγ signal.

We noticed that PTPN1_S270/Y271 and PTPN2_S268/Y269 are homologues residues, implying that they might exert their regulatory effects through similar mechanisms. We selected PTPN1_S270 and PTPN2_S268 as representatives and confirmed that their respective targeting sgRNAs generated the expected mutations with minimal bystander editing (Supplemental Information). To verify the function of these dominant mutations, we separately overexpressed the WT cDNA and the corresponding mutants in A375 cells. Introduction of PTPN1_S270P or PTPN2_S268P variant decreased the expression of PTPN1 or PTPN2, respectively. This, in turn, resulted in an upregulation of PD-L1 in both total and membrane protein levels (Figure S3I). To comprehensively investigate the regulation of these two endogenous mutations in A375 cells, we focused on examining typical proteins involved in IFNγ signaling pathway. Both mutations increased the protein levels of JAK2, subsequently leading to a significant upregulation in pSTAT1 levels without changing the overall abundance of STAT1 protein (Figure 3J; Figure S3J). These results suggest that PTPN1_S270P and PTPN2_S268P activate the IFNγ pathway by reducing the abundance of each respective protein, ultimately resulting in an increased pSTAT1 level and, consequently, an upregulation of PD-L1 expression. Similarly, we found that multiple mutations identified in PTPN1 and PTPN2 also led to a reduction in their own protein levels and an increase in the total and membrane PD-L1 abundance (Figure S3K; Figure 2D). For these loss-of-function (LOF) mutations, their subsequent effects were consistent with the outcomes of knocking out PTPN1 or PTPN2 using the CRISPR/Cas9 system (Figure S3L).

We have summarized the critical S/T/Y residues within the SH2-B and PTP family proteins for depicting their regulatory effects on the canonical IFNγ pathway (Figure 3K). Based on the screening and validation results, in combination with prior related studies, we have created a gene network diagram outlining PD-L1 modulation at the single amino acid level (Figure S4). The rich information of functional residues contributes to a better understanding of the roles played by these corresponding proteins and provides initial insights for refining the PD-L1 regulatory network from a single residue perspective.

### Genome-wide mapping of critical residues modulating HLA-I expression using S/T/Y library

Tumor cells can employ various strategies to evade immune surveillance. In addition to increase the expression of immune checkpoint ligands, defects in MHC-I-mediated antigen processing and presentation (APP) can directly hinder the tumor recognition of CD8+ T cells and restrain their activation and proliferation (Jhunjhunwala et al., 2021). Genetic mutations in essential genes involved in MHC-I APP have been implicated in tumor progression and the development of resistance to ICB therapy (Gettinger et al., 2017; Shin *et al*., 2017). Therefore, beyond interpreting the regulation of PD-L1 pathway, it is also crucial to understand the regulatory mechanisms of MHC-I in tumor cells.

We thus performed an additional S/T/Y library screen to investigate the functional residues that modulate HLA-I expression in A375 cells. Using the pan-human HLA-I-specific antibody W6/32 for protein staining, we observed a relatively high level of surface HLA-I expression in A375 cells without IFNγ stimulation, which enables to identify both positive and negative regulators of HLA-I expression in this context (Figure S5A). Consequently, we conducted the library screen for HLA-I regulators in A375-ABEmax cells in the absence of IFNγ. Through the same procedure of FACS enrichment and data analysis as described for the PD-L1 screen (Figure 1A; Figure S5B-C), we identified numerous sites within genes related to APP that were prominent in HLA^low^ cells (Figure 4A). These regulators included multiple allelic variants of HLA, the TAP binding protein TAPBP (tapasin), antigen transporters TAP1 and TAP2, and the component of MHC-I complex, B2M. In the HLA^high^ group, we also observed novel sites enriched on several negative regulators of HLA-I, including SUSD6_Y177 and WWP2_Y704, whose coding genes were recently reported to form an HLA-I inhibitory axis (SUSD6/TMEM127/WWP2) for cancer immune evasion (Chen et al., 2023).

**Figure 4.**
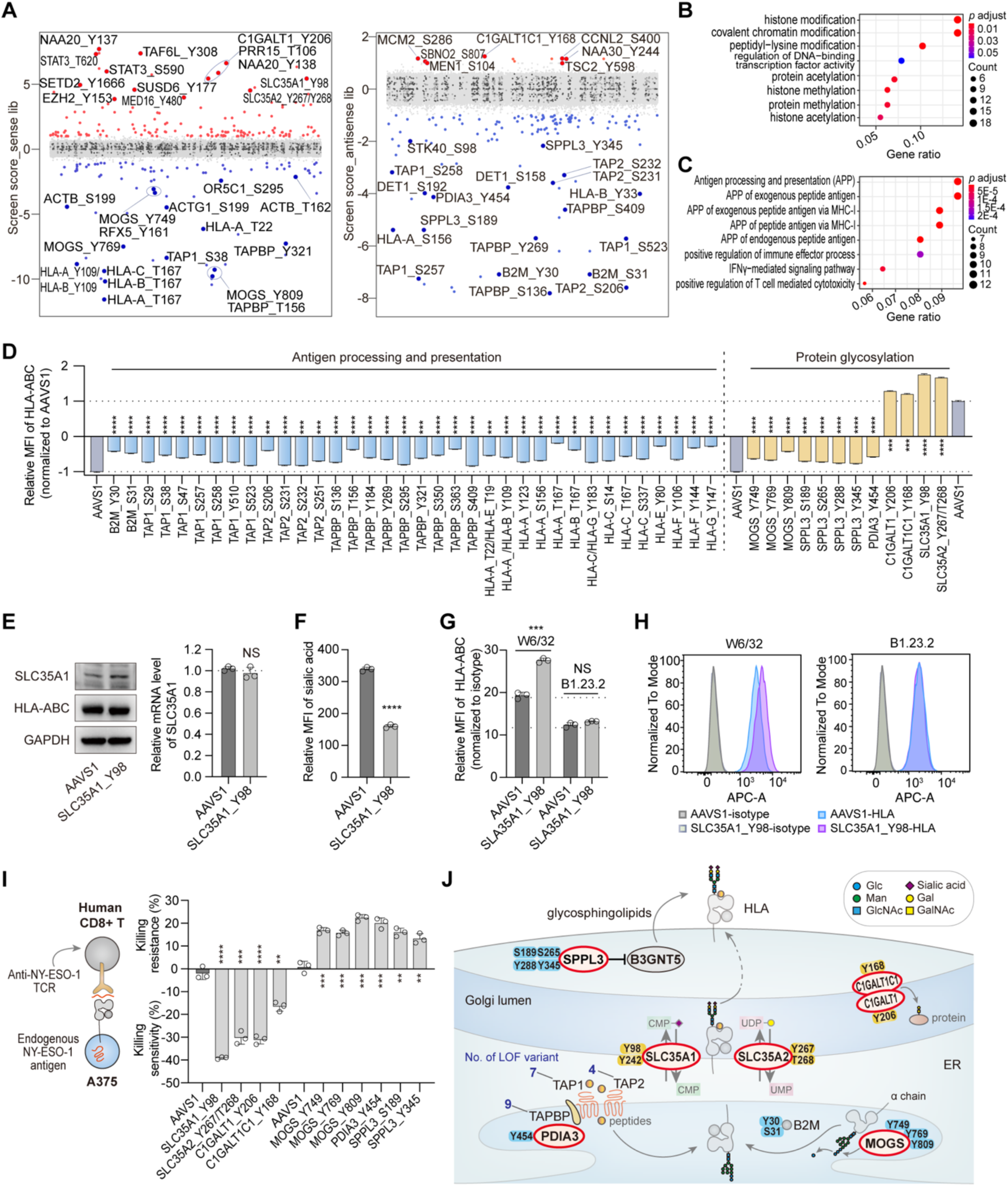
ABE-based screens identify functional S/T/Y residues modulating HLA-I expression in genome-wide. (A) Significant S/T/Y residues enriched from the sense library (left) and antisense library (right) that upregulate or downregulate HLA-I expression in A375 cells without IFNγ stimulation. The screening procedure is the same as Figure 1A. Positively or negatively enriched sites were selected by screen score > 1 or < -1. (B-C) GO enrichment analysis of related genes with identified mutations leading to HLA-I upregulation (B) and downregulation (C) in the absence of IFNγ. (D) Individual validations of representative sites related to APP and protein glycosylation in A375 cells in the absence of IFNγ by flow cytometry analysis. The method to generate relative MFI of HLA-ABC and the statistics are the same as that shows in Figure 2C-D. (E) Protein expression levels of SLC35A1 and HLA-ABC (left) and relative mRNA expression levels of *SLC35A1* (right) in A375 cells infected with sgRNA targeting *AAVS1* and SLC35A1_Y98. The mRNA level of each sample was quantified by real-time qPCR and normalized by *GAPDH*. For each mutant cells, the indicated relative mRNA level was normalized to that of *AAVS1*-targeting control cells. The data was presented as the mean ± SD (n=3). *P* values were calculated using Student’s *t* test, NS, not significant. (F) Relative MFI of surface sialic acid of A375 cells infected with sgRNA targeting *AAVS1* and SLC35A1_Y98 by flow cytometry analysis. The data was presented as the mean ± SD (n=3) and normalized to that of isotype. *P* values were calculated using Student’s *t* test, \*\*\**P* < 0.001. (G-H) Relative MFI (G) and flow cytometry histograms (H) of surface HLA-ABC of A375 cells infected with sgRNA targeting *AAVS1* and SLC35A1_Y98 using different HLA-I-specific antibodies for staining. In Figure 4G, the data was presented as the mean ± SD (n=3) and normalized to that of isotype. *P* values were calculated using Student’s *t* test, \*\*\**P* < 0.001, NS, not significant. (I) Killing resistance and sensitivity of A375 cells infected with sgRNAs targeting residues on glycosylation-related genes to expanded NY-ESO-1 CD8+ T cells. The data was presented as the mean ± SD (n=3). *P* values were calculated using two-tailed Student’s *t* test, \*\*\**P* < 0.001, \*\*\**P* < 0.001, \*\*\*\**P* < 0.0001. (J) Schematic diagram of the HLA-I regulatory network focused on identified residues on representative APP and glycosylation-related genes. See also Figure S5-6 and Table S5-8.

Upon integrating all the relevant genes identified through the screen, a GO analysis about the biological process revealed that some similar terms to those from the PD-L1 screen were among the top-ranked in HLA^high^ group. These terms included processes related to histone modification, covalent chromatin modification, and peptidyl-lysine modification, highlighting the general regulatory influence of genes on both PD-L1 and HLA-I (Figure 4B). In contrast, in the HLA^low^ group, multiple terms related to antigen processing and presentation were highly enriched (Figure 4C). STRING PPI network analysis of top-ranked regulators further revealed genes involved in antigen processing and presentation, immune response, and cellular protein modification process (Figure S5D). Of note, we identified several nodes connecting multiple networks, such as HLA-A, B2M, indicating their central roles in regulating the expression of each respective protein.

### Interpretation of novel residues regulating antigen recognition and presentation

Compared with the candidate sites identified in the PD-L1 screens, the HLA-I screens revealed a multitude of unique variants with unknown functions that were enriched in both high and low directions, not limited to sites within APP-related genes. To further assess their impact on surface HLA-I expression, we conducted a large-scale verification of candidate sites enriched on various regulatory pathways (Figure 4D; Figure S6A).

In the HLA-I screens, a dominant category of functional sites enriched in APP-related genes. Nearly all relevant residues were subjected to individual validation, all of which were verified to significantly reduce surface HLA-I expression upon targeted mutation (Figure 4D). For the gene *B2M*, two noteworthy sites, Y30 and S31 located in its Ig-like C1-type domain (Figure S6B), showed significant phenotypic effects. Targeting each of these two sites led to a double mutation, Y30H and S31P, which led to reduced levels of both membrane and total HLA, while B2M expression remained unaltered (Figure S6C-D). As for the *TAP1* and *TAP2* genes, multiple hits within TAP1 were mainly localized in its N-terminal domain and ABC transmembrane type-1 domain, and functional residues of TAP2 converged on its ABC transmembrane type-1 domain (Figure S6E). Unlike the mutants of B2M, several top-ranked mutations of TAP1 not only disrupted its own expression but also further reduced HLA expression at both total and membrane protein levels (Figure S6F). As for *TAPBP* (Figure S6E), we investigated eight mutants with significant phenotype of reduced surface HLA levels, among which five mutations slightly reduced the overall HLA expression and three had no significant effect on total HLA levels (Figure S6G).

In particular, we identified a significant number of mutations in glycosylation-related genes that were enriched in both HLA^high^ and HLA^low^ groups. Among these, the SLC35A1_Y98 mutation led to a dramatic increase in surface HLA expression (Figure 4D). SLC35A1 is a membrane-bound transporter located in the Golgi apparatus responsible for transferring CMP-sialic acid from the cytosol into the Golgi apparatus. This process facilitates the sialylation of proteins by various sialyltransferases. Importantly, the Y98C mutation did not interfere with the expression of SLC35A1 at both the protein and mRNA levels (Figure 4E). The residue Y98 is involved in the binding of CMP and CMP-sialic acid and is essential for optimal transport competence, as confirmed by previous *in vitro* studies (Nji et al., 2019). We thus examined the sialic acid level on the cell membrane and found that SLC35A1_Y98C mutation significantly impaired the transport of sialic acid (Figure 4F). To further explain the relevance between SLC35A1 mutation and HLA abundance, we assessed the overall HLA level in SLC35A1_Y98 mutant cells. Intriguingly, we found that the mutation increased the median fluorescence intensity (MFI) of surface HLA without changing overall HLA expression (Figure 4D-E). Prior study has reported that sialic acid residues on glycosphingolipid (GSL), synthesized by B3GNT5, is involved in shielding critical epitopes of HLA-I molecules on the cell surface, thus diminishing their interactions with several immune cell receptors and decreasing CD8+ T cell responses (Jongsma et al., 2021). Therefore, we investigated whether sialic acid modification affected the accessibility of the HLA-I epitope recognized by the W6/32 antibody, which is commonly used for HLA-I staining in multiple genetic screens (Dersh et al., 2021; Jongsma et al., 2021; Sparbier et al., 2023). Using another HLA-I antibody, B1.23.2, with different recognition epitope, we detected no significant change in surface HLA-I levels after mutating SLC35A1_Y98, indicating that glycosylation modifications are indeed crucial for the epitope recognition of HLA-I specific antibodies, as well as for immune cell receptors (Figure 4G-H).

Similarly, several sites were identified in additional glycosylation-related genes. These genes included negative regulators, such as another nucleotide sugar transporter called SLC35A2 responsible for transporting UDP-galactose, as well as the glycosyltransferase C1GALT1 and its chaperone C1GALT1C1. On the positive regulatory side, there were genes like *SPPL3*, which negatively regulates B3GNT5 expression, thereby controlling GSL synthesis (Jongsma et al., 2021). Additionally, MOGS (α-glucosidase I), which encodes the first enzyme responsible for trimming N-glycans in the endoplasmic reticulum (ER) (Varki et al., 2022), and PDIA3 (ERp57), which is involved in the general glycoprotein folding process within the ER and is required for optimal tapasin activity (Wearsch and Cresswell, 2007), also had identified sites. While most of the residues identified in these glycosylation-related genes exhibited no effect on the overall HLA-I expression, they significantly influenced surface HLA-I levels (Figure S6H). Therefore, it is likely that these residues can affect the glycosylation of various regulators or perturb the structural stability of glycoproteins involved in APP process.

To validate the significance of these glycosylation-related residues in immunosurveillance, we examined the susceptibility of these mutants to CD8+ T cell-driven cytotoxicity. We generated each mutation in A375-ABEmax cells, which endogenously express HLA-A2 and NY-ESO-1 antigen. Human CD8+ T cells transduced with an HLA-A2-restricted T cell receptor (TCR) specific for the NY-ESO-1 antigen (Robbins et al., 2008) were co-cultured with the mutant cells. We found that the HLA^high^ variants were more sensitive to T cell-driven killing, with SLC35A1_Y98 being an example. In contrast, the HLA^low^ mutations conferred significant resistance to T cell killing, thus subverting T-cell-mediated immunosurveillance (Figure 4I). These novel residues have been summarized for their functional roles in different glycosylation processes (Figure 4J), which can alter the glycosylation of related regulatory genes or impact the quality control machinery of glycoprotein involved in the assembly of HLA-I molecules, consequently affecting the recognition of tumor cells by T cells.

### Integrated analysis for potential co-regulators of surface PD-L1 and HLA-I

The above analyses drew a comprehensive map of regulators for surface PD-L1 and HLA-I at the residue level. However, in the *in vivo* tumor microenvironment, various factors collectively influence the fate of tumor cells. To gain a deep understanding of the co-regulators of PD-L1 and HLA-I, two of the principal factors for immunotherapy, we conducted a comparison of candidates from the HLA-I and PD-L1 screens in the presence and absence of IFNγ stimulation (Figure 5A). This analysis revealed five mutants that upregulated HLA-I expression and downregulate PD-L1 expression in the presence of IFNγ, including Y137 and Y138 of the N-terminal acetyltransferase NAA20. These mutants are likely to function as positive regulators of antitumor immunity. Conversely, one mutant, MAPK3_Y333, was found to downregulate HLA-I expression and upregulate PD-L1 expression, indicating its potential role in promoting tumor evasion. Additionally, we identified 13 mutants that concurrently upregulated HLA-I and PD-L1 expressions, including six hits that increased PD-L1 levels upon IFNγ treatment, such as EZH2_Y153, EED_Y308, and SETD2_Y1666, all of which are involved in epigenetic modulations.

**Figure 5.**
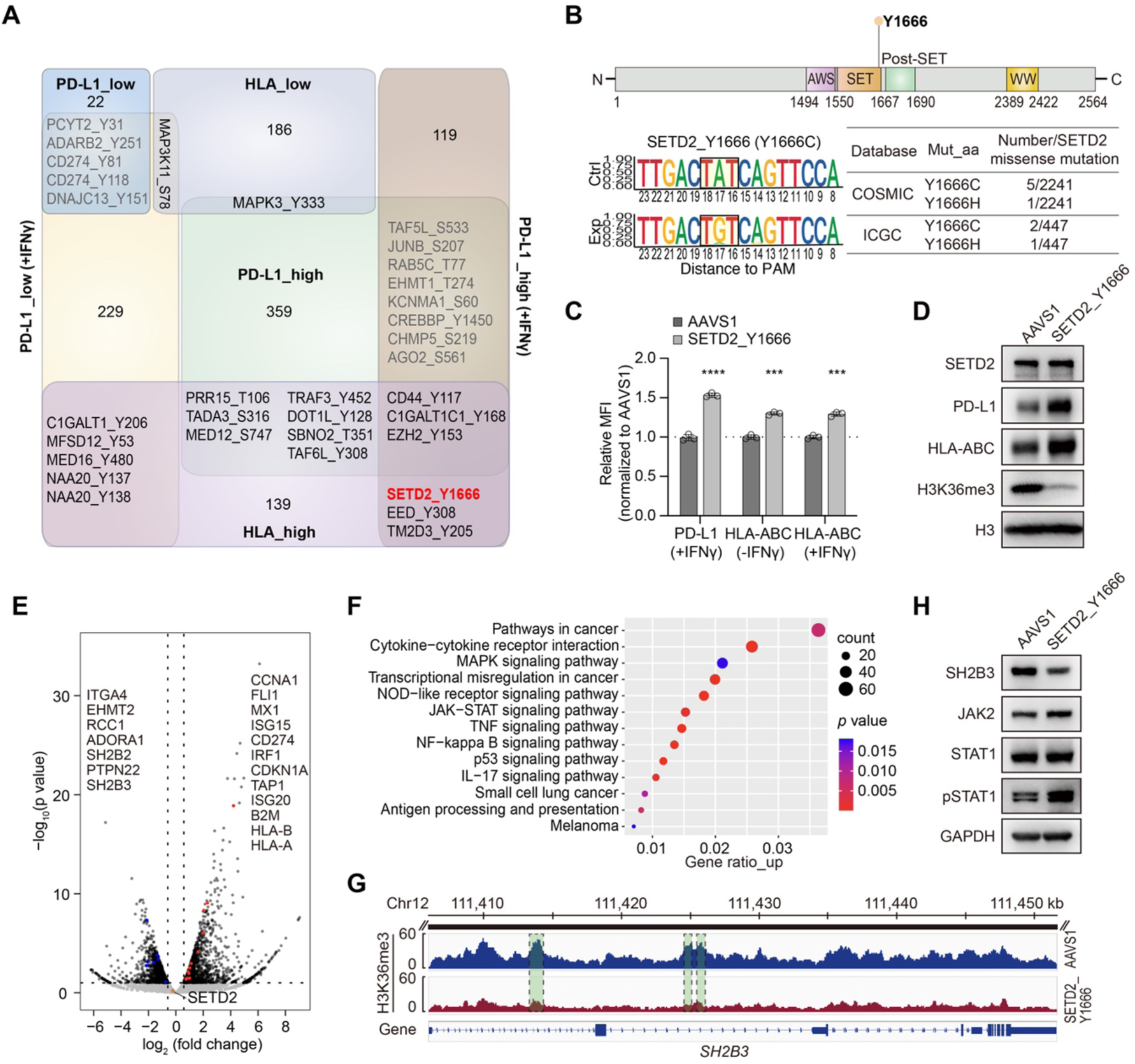
Interpretation of functional residues co-regulating surface PD-L1 and HLA-I. (A) Comparison of S/T/Y residues identified from PD-L1 screens and HLA-I screens using venn diagram. (B) General information of SETD2_Y1666. The upper structure schematic indicates the location of Y1666 residue on SETD2 protein. The lower figures (left) indicate the editing outcomes of sgRNA targeting SETD2_Y1666 by NGS analysis. The lower table (right) indicates the information of clinical relevance of SETD2_Y1666. (C) Relative MFI of surface PD-L1 and HLA-I of A375 cells infected with sgRNA targeting *AAVS1* and SETD2_Y1666 with different IFNγ treatment. The method to generate relative MFI of PD-L1 or HLA-I and the statistics are the same as that shows in Figure 2C-D. (D) Protein expression levels of SETD2, PD-L1, HLA-ABC and H3K36me3 in A375 cells infected with respective sgRNA targeting *AAVS1* and SETD2_Y1666. (E) Volcano plots showing the DEGs between SETD2_Y1666-targeted A375 mutant cells and *AAVS1*-targeted A375 control cells. The represented genes are listed. (F) Representative KEGG pathway analysis of upregulated DEGs in SETD2_Y1666-targeted A375 mutant cells compared with the *AAVS1*-targeted control. The DEGs were selected using the threshold of FC > 1.5 and *p* value < 0.1 according to the RNA-seq data. (G) ChIP-seq tracks for H3K36me3 at *SH2B3* gene locus between SETD2_Y1666-targeted A375 mutant cells and *AAVS1*-targeted A375 control cells. (H) IB analysis of SH2B3 and typical JAK/STAT signaling components in A375 cells infected with respective sgRNA targeting *AAVS1* and SETD2_Y1666. See also Figure S7 and Table S7-8.

To explore the regulatory mechanisms of these novel co-regulators, we focused on the functional investigation of the category with the largest number of mutants, which increased the expression level of both PD-L1 and HLA-I. Among them, SETD2_Y1666, as well as the corresponding coding gene, stood out as a novel regulator, whose relevance with PD-L1 or HLA-I has not yet been reported. SETD2 is the primary histone methyltransferase responsible for catalyzing H3K36me3, representing a marker of transcriptional activation. SETD2 is associated with diverse biological functions, such as maintenance of genomic stability (Park et al., 2016), antiviral immune response (Chen et al., 2017), and restriction of tumor metastasis (Yuan et al., 2020). SETD2 mutations are prevalent in various human tumors and are reported to be associated with tumor progression, including glioma, clear cell renal cell carcinoma, leukemia, and prostate cancer (Armenia et al., 2018; Cancer Genome Atlas Research, 2013; Fontebasso et al., 2013; Zhu et al., 2014). We found that Y1666 is in the SET domain of SETD2, which is the catalytic domain mediating the H3K36me3-specific methyltransferase activity (Sun et al., 2005). SETD2_Y1666 targeted by ABEmax could generate the Y1666C mutation, a reported mutation from both COSMIC (Catalogue of Somatic Mutations in Cancer) and ICGC database (Figure 5B). We found that Y1666C didn’t change the expression of SETD2 at both the mRNA and protein levels, but it significantly increased the total and membrane protein levels of PD-L1 and HLA-I upon IFNγ exposure (Figure 5C-D). Meanwhile, the expression level of H3K36me3 was markedly decreased, suggesting that the Y1666C mutation disrupted the catalytic activity of SETD2 without affecting its own protein expression (Figure 5D).

To elucidate the mechanisms of PD-L1 and HLA-I regulation by SETD2_Y1666, we performed RNA-seq and H3K27me3 ChIP-seq analysis for SETD2_Y1666C mutant cells and control cells with IFNγ stimulation, gaining insight into the potential targets of SETD2. We analyzed the differential expressed genes (DEGs) from the RNA-seq data, and identified numerous representative upregulated DEGs in the mutant cells, as exemplified by *CD274*, *IRF1*, *TAP1*, *B2M*, *HLA-A*, *HLA-B* and *HLA-C*, all of which are directly associated with PD-L1 and HLA-I expressions (Figure 5E). By analyzing the enriched KEGG pathways of upregulated genes, we found dominant terms, including cytokine-cytokine receptor interactions, transcriptional misregulation in cancer, NF-κB signaling pathway, JAK-STAT signaling pathway, and antigen processing and presentation (Figure 5F). We further referred to the ChIP-seq data to search for the methylated targets of SETD2 and found genes with a significant reduction in H3K36me3 signal, such as *RCC1* (Figure S7A), which was reported to enhance PD-L1 expression and improve the ICB sensitivity after gene knockdown (Zeng et al., 2021). *RCC1* was also downregulated in the RNA-seq analysis (Figure 5E), indicating that SETD2_Y1666 mutation could decrease the H3K36me3 modification of *RCC1*, thus upregulating PD-L1 expression. Interestingly, multiple gene body regions of *SH2B3* exhibited a remarkable lower H3K36me3 signal (Figure 5G), and SH2B3 appeared to be downregulated upon SETD2_Y1666 mutation (Figure 5E). Considering the negative regulation of SH2B3 on the JAK-STAT signaling pathway (Figure 3F-G) and the detected enrichment of JAK-STAT signaling in SETD2_Y1666C mutant cells (Figure 5F), we further investigated the effects on this pathway when Y1666 was mutated. We found that SETD2_Y1666C conferred a significant reduction in SH2B3 expression, along with higher expression of JAK2 and pSTAT1 (Figure 5H; Figure S7B), which correlated with the effects of SH2B3 mutants. We also detected upregulation of IFNγ responsive genes such as *IRF1*, some interferon stimulated genes including *ISG15*, *ISG20*, and *MX1* (Figure 5E; Figure S7B), which are associated with the upregulation of PD-L1 and HLA-I (Burks et al., 2015; Garcia-Diaz et al., 2017). The above analysis revealed that the Y1666 mutation in the SET domain of SETD2 could boost JAK-STAT signaling pathway, thus increasing PD-L1 expression and antigen processing and presentation.

Besides SETD2_Y1666, we also investigated another category of mutants enriched in HLA^high^ and PD-L1^low^ group, represented by two clinical mutations NAA20_Y137 and Y138 (Figure 6A; Figure S7C). Targeting either of them with ABEmax could generate Y137C and Y138C co-mutations, we thus used NAA20_Y137 as a representative (Figure S7D). We found that targeting NAA20_Y137 didn’t affect the protein level of NAA20 but resulted in PD-L1 reduction and HLA-I upregulation in both membrane and total protein levels with IFNγ treatment (Figure S7E-F). Overexpressing mutated cDNAs of NAA20_Y137C, Y138C, and Y137C/Y138C in A375 cells indicated that all three mutants contributed to the modulation of PD-L1 and HLA-I expressions (Figure S7G). RNA-seq analysis further revealed that targeting NAA20_Y137 can lead to downregulation of several dominant KEGG terms, including MAPK, PI3K-Akt, TNF, and NF-κB signaling pathways, along with upregulated terms, such as DNA replication and APP (Figure S7H-I). Given that the two residues are located in the N-acetyltransferase domain of NAA20 and involved in mediating the interaction between NAA20 and its catalytic substrate (Deng et al., 2020) (Figure S7C), we hypothesized that the mutations may disrupt NAA20’s N-acetyltransferase activity and affect the acetylation of its substrate, thus co-regulating PD-L1 and HLA-I expressions.

**Figure 6.**
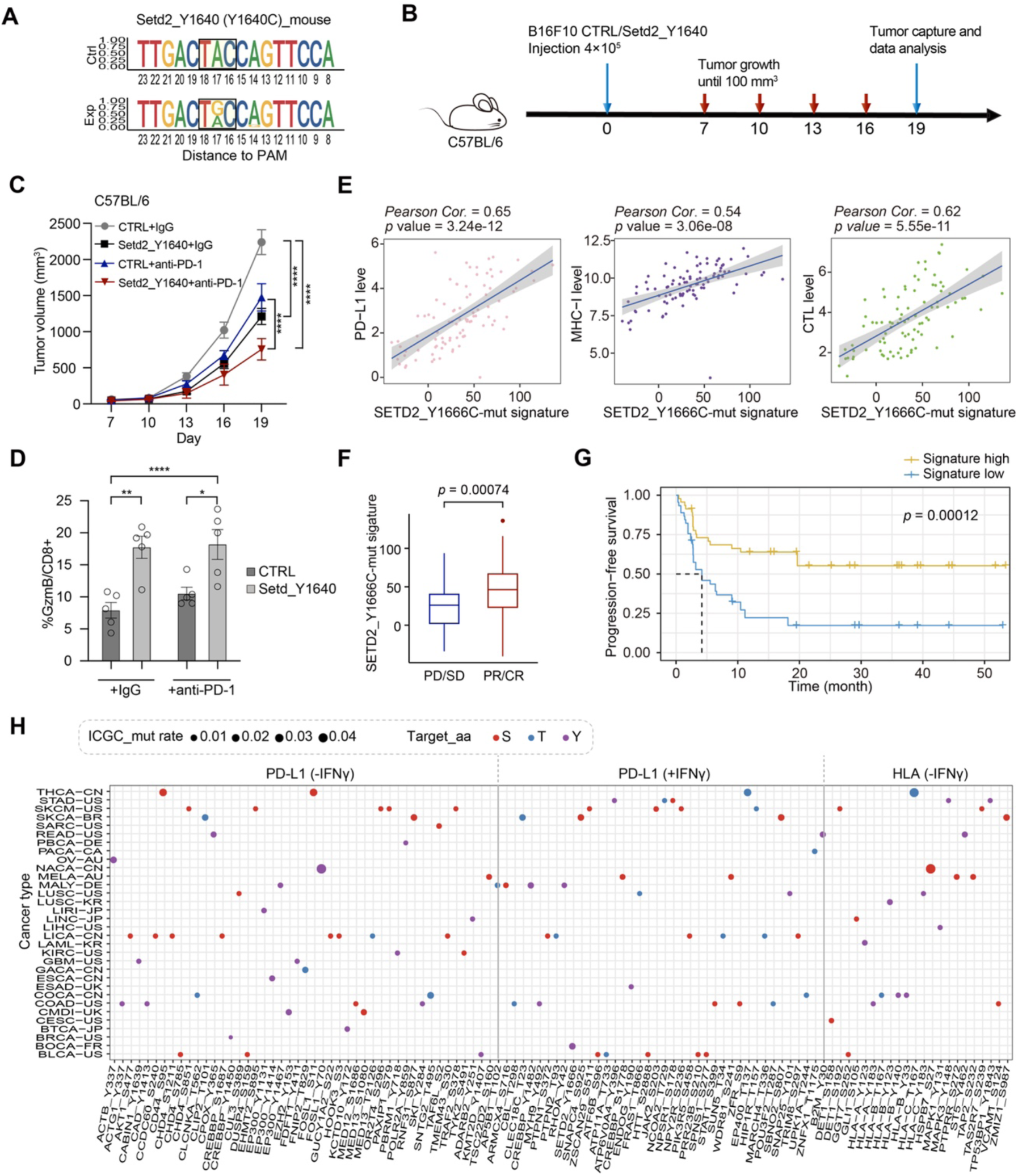
Clinically relevant mutation SETD2_Y1666/Setd2_Y1640 contributes to an improved response to ICB therapy *in vivo*. (A) Editing outcomes of sgRNA targeting Setd2_Y1640 by NGS analysis. Ctrl and Exp respectively indicates the WT and mutated sequence in B16F10 cells. (B) A schematic view of implanting B16F10 mutant cells and CTRL cells to C57BL/6 mice and the following treatment of PD-1 mAb or IgG isotype control (IgG2a). (C) Longitudinal tumor size of the indicated B16F10 tumors in C57BL/6 mice treated by control IgG or ICB. The data was presented as the mean ± S.E.M. (n = 5 mice/group) for each group at each time point. *P* values were calculated using Two-way ANOVA with Benjamini-Hochberg adjustment for multiple testing, \*\*\*\**P* < 0.0001. (D) Quantification of GzmB represented as percentage on CD8+ TILs in B16F10 tumors harvested from C57BL/6 mice after indicated treatments. The data was presented as the mean ± SD (n = 5 mice/group) for each group at each time point. *P* values were calculated using Student’s *t* test, \**P* < 0.05, \*\**P* < 0.01, \*\*\*\**P* < 0.0001. (E-G) Correlation between SETD2_Y1666-mutation signature and PD-L1 expression, MHC-I expression, intratumoral CTL infiltration (E), ICB response (F), and overall survival and progression-free survival (G) in patients treated by anti-PD-1 in the Gide *et al*. study (Gide *et al*., 2019) in melanoma. PD: progressive disease, SD: stable disease, PR: partial response; CR: complete response. *P* value was respectively calculated by two-tailed Student’s *t* test (F) and log-rank test (G). (H) Schematic of representative residues identified from PD-L1 and HLA-I screens with clinical relevance according to ICGC database. X axis indicates functional residues regulating PD-L1 or HLA-I from the ABE screens. Y axis indicates different cancer types defined in ICGC database. The dot size represents the detected missense mutation rate of each indicated residue. See also Figure S8-9 and Table S8.

### Functional clinical mutations promote cancer immunotherapy *in vivo*

Considering the regulatory impact of these clinically relevant mutations on both PD-L1 and HLA-I, we intended to dissect their potential effects on tumor progenesis and response to ICB treatment *in vivo*. We thus created the homogenous mutations using ABEmax system in a mouse melanoma cell line, B16F10, corresponding to the human mutations SETD2_Y1666C. The sgRNA targeting Setd2_Y1640 were infected into B16F10-ABEmax cell line, which resulted in similar editing patterns as observed in A375 cells (Figure 6A). Subsequently, we separately injected the Setd2_Y1640-targeted B16F10 cells into the immune-competent C57BL/6 mice, as well as negative control samples infected with an sgRNA targeting the safe-harbor locus, to establish B16F10 melanoma tumors (Figure 6B). As expected, we observed a significant reduction in tumor growth in Setd2_Y1640-targeted mice, and the combination of anti-PD-1 treatment further inhibited tumor progression (Figure 6C). Meanwhile, we observed a consistent tumor growth pattern between the mutant group and the control in the immune-deficient BALB/C nude mice (Figure S8A), indicating that these two mutations contribute to tumor suppression only through reshaping the immune microenvironment.

To further investigate the impact of Setd2_Y1640 mutation and its combination with ICB treatment on TME, we analyzed infiltrated immune cells in B16F10 tumor-bearing C57BL/6 mice. In Setd2_Y1640 mutant group, we detected an increased expression of the T cell activation marker Granzyme B (GzmB) on infiltrated CD8+ T cells compared to the control, and the combination of ICB treatment further strikingly elevated the percentage and activation of CD8+ T cells (Figure 6D). These results indicated that the mutation might reshape the TME through the activation of representative signaling pathways, including NF-κB and JAK-STAT, which in turn upregulated PD-L1 and HLA-I expression. This enhances the cytotoxicity of tumor infiltrating CD8+ T cells and improves the efficacy of anti-PD-1/PD-L1 blockade therapy *in vivo*.

After confirming the effects on immune response in the mouse model, we next attempted to analyze the correlation between genetic mutation-derived functional deficiency and the response to immunotherapy in published ICB treatment cohorts. We first derived the gene expression signature of the SETD2_Y1666C mutation based on its RNA-seq results, as described in a previous study (Gu et al., 2021). Referring to 91 RNA-seq samples from 54 patients in a melanoma cohort treated with anti-PD-1 (Gide et al., 2019), we confirmed that the SETD2_Y1666C-mutation signature was positively correlated with tumor PD-L1, MHC-I, and cytotoxic T-cell infiltration (Figure 7E). Further analysis revealed that patients responding to ICB therapy (partial response and complete response: PR/CR) exhibited higher SETD2_Y1666C-mutation signature compared with non-response groups (progressive disease and stable disease: PD/SD) (Figure 7F), and the mutation signature also showed a positive correlation with progression-free survival (Figure 7G). Interestingly, recent studies also found that patients with different cancer types that harboring SETD2 deleterious mutations showed improved response to ICB therapy (He et al., 2023; Lu et al., 2021). Collectively, these findings firstly demonstrated the mechanisms of SETD2_Y1666C mutation in modulating immune surveillance and further supported the notion that the mutation is relevant to a better response to ICB treatment in clinical trials. Due to the high mutation rate of SETD2 in various cancer types, SETD2 may serve as a biomarker for ICB treatment and a large population of patients may benefit from immunotherapy.

We also created the homologous variants Naa20_Y137C/Y138C using sgRNA targeting Naa20_Y137 site in B16F10 cells (Figure S8B) and assessed its impact on immune response *in vivo* (Figure 6B). Similar as SETD2_Y1666, a significant reduction in tumor growth was observed in Naa20_Y137C/Y138C mutant group and anti-PD-1 treatment further restrained tumor progression (Figure S8C-D). In-depth analysis revealed that Naa20_Y137C/Y138C mutation led to a significant increase in the percentage of infiltrated CD8+ T cells expressing GzmB, which was further elevated in the ICB combination group (Figure S8E).

In addition to the clinical mutations SETD2_Y1666C and NAA20_Y137C/Y138C described above, we sought to investigate the clinical relevance of all selected mutants identified in the screens. Referring to different sequencing data from cancer patients, including ICGC and COSMIC, we found 168 sites with detected mutations across 35 tumor types in ICGC (Figure 6H; Figure S9A), and more than 300 sites recorded in COSMIC (Figure S9B-D). Overall, nearly 40% (416/1083) of the identified residues from the three screens were clinically observed in these databases, providing a rich resource of potential pathogenic mutations, especially those linked to cancer. Furthermore, this information offers guidance on the efficacy of ICB for patients harboring these mutations.

## DISCUSSION

In this report, we conducted a large-scale sgRNA library screen using the ABE system to identify functional genetic variants that modulate the expression of two crucial determinants in cancer immune response: PD-L1 and HLA-I. These factors play a pivotal role in the effectiveness of ICB therapy, particularly in the context of PD-1/PD-L1 blockade. We employed a specialized library targeting 584,377 sites across the genome, encompassing all designable residues of serine, threonine, and tyrosine. Through this approach, we successfully identified over 1,000 novel sites associated with the upregulation or downregulation of PD-L1 or HLA-I expression, using stringent criteria. These identified residues are enriched in several critical immune-related pathways, such as chromatin remodeling, histone modification, JAK/STAT signaling, and antigen processing and presentation. This comprehensive mapping provides valuable insights into the regulation of cancer immune responses at both the amino acid and base levels for the first time.

To systematically identify critical sites involved in immune response regulation, we initiated our investigation by focusing on one of the key PTMs, phosphorylation. Phosphorylation is known to play a crucial role in signaling transduction and the regulation of gene expression. Our screens identified numerous residues on well-known genes as well as novel genes associated with IFNγ-induced JAK/STAT signaling, including IFNγ receptors, JAK kinases, STATs, and proteins from SH2-B family and PTP family. Among these sites, there was a significant enrichment of well-known phosphorylation sites, including STAT1_Y106 and Y701, JAK1_Y806 and Y830, and JAK2_Y1007 and Y1008. Additionally, we uncovered several predicted phosphorylation sites, exemplified by multiple sequential sites that were concentrated within the SH2 domain of STAT1 and SH2B3. Mutations at these phosphorylation sites have the potential to deactivate the target genes and result in the disruption of phosphorylation events, ultimately leading to the downregulation of PD-L1 or HLA-I. Importantly, our screens were not limited to investigating phosphorylation sites. Amino acid substitutions can lead to either decreased, increased, or unchanged protein levels, resulting in gene inactivation or augmentation. Thus, different from canonical CRISPR/Cas9 screens, which primarily focus on gene-level dysfunction, base editing-based screens allow for both LOF and GOF perturbations in a single screen. For instance, we found that for the positive regulators of JAK/STAT signaling, such as STAT1 and STAT3, our screens identified mutations that either downregulated or upregulated gene expressions. We found that the majority mutations downregulate gene expressions, which may affect mRNA or protein stability, including known phosphorylation sites such as STAT1_Y106 and Y701. Additionally, certain number of mutations did not alter the expression level of the targeted genes but could affect DNA binding capacity, such as STAT1_S462, disrupt protein-protein interactions, such as mutations on SH2-B adaptor proteins and CBL, or impair enzymatic catalytic activity, such as SETD2_Y1666 and NAA20_Y137/Y138. These functional sites unveiled novel and comprehensive mechanisms of cancer immune response regulation, which cannot be fully explored through gene-level screens alone.

While previous studies have investigated the regulation of PD-L1 or HLA-I through separate CRISPR screens, the coordinated regulation of both PD-L1 and HLA-I has not been systematically explored, especially at the residue level. In this report, based on functional residue screens for PD-L1 and HLA-I regulation, we identified numerous sites that specifically modulate each factor. Furthermore, we highlighted novel residues that simultaneously modulate PD-L1 and HLA-I expression and delved into their *in vivo* functions. We focused on two such variants, SETD2_Y1666 and NAA20_Y137/Y138, which upregulated HLA-I expression while affecting PD-L1 levels in opposite directions. Notably, the functional roles of these genes in PD-L1 or HLA-I regulation have not been previously reported. We discovered that mutations in these genes significantly impaired the interaction between enzymes and their substrates or the catalytic activity of the enzymes, without affecting the protein expression levels. Intriguingly, both variants promoted immune responses and enhanced the efficacy of anti-PD-1 immunotherapy, with HLA-I upregulation likely playing a leading role in these scenarios. For SETD2_Y1666, the upregulation of HLA-I-dependent antigen presentation appeared to counterbalance the adverse effect of PD-L1-mediated immune evasion, reshaping the tumor immune microenvironment to favor anti-PD-1/PD-L1 immunotherapy. Additionally, SETD2-dependent PD-L1 induction could also enhance the effectiveness of anti-PD-1 blockade to restore suppressed antitumor immunity. In support of our view, previous studies also reported that deficiencies in negative regulators of PD-L1, such as ADORA1 (Liu et al., 2020), UROD (Suresh et al., 2020), and USP8 (a negative regulator of both PD-L1 and HLA-I) (Xiong et al., 2022), can enhance the therapeutic effects of anti-PD-1/PD-L1 immunotherapy *in vivo*.

Consistence with a previous base editing screen for IFNγ signaling regulators (Coelho et al., 2023), many of the functional mutations identified in our screens have clinical precedence, as supported by data from the ICGC and COSMIC databases. This suggests the prevalence of cancer immunoediting and highlights the clinical significance of these mutations. For both clinically characterized and uncharacterized mutations, our multidimensional screens unveiled their potential impacts on cancer development and progression. Additionally, our dataset provides clinically relevant biomarkers for predicting immune response and resistance to ICB treatment, while also suggesting novel strategies for combinational immunotherapy. Moreover, multiple CRISPR/Cas9 screens have identified a series of PD-L1 or MHC-I regulators that can serve as druggable targets, such as CMTM6 (Burr et al., 2017; Tu et al., 2019), EZH2 (Burr et al., 2019; Dersh et al., 2021), TRAF3 (Gu et al., 2021), highlighting the importance of combination therapy with ICB. The base-level screens presented in this study not only revealed the importance of single residues but also identified several novel genes, including *FECH*, *TAF5L*, *TAF6L*, *CHMP5*, *NAA20*, and *SETD2*, further enriching the resource of potential therapeutic targets for combination ICB therapy. Importantly, the base-level information provides mechanistic insights that can guide the development of novel drugs.

To sum up, our study provides a comprehensive resource of functional residues involved in the regulation of PD-L1 and HLA-I, shedding light on the understanding of human immune responses at the base level. This initial step in mapping the regulatory residues involved in immunosurveillance can be further complemented by investigating other PTMs, such as ubiquitylation, and by employing other gene editing tools, including prime editors (Anzalone et al., 2019; Chen et al., 2021a; Nelson et al., 2022) or PAMless Cas9-based base editors (Walton et al., 2020), to further expand the coverage of amino acids.

## ACKNOWLEDGMENTS

We acknowledge National Center for Protein Sciences (Beijing) of Peking University for assistance with fluorescence-activated cell sorting and analysis, particularly Dr. Jia Luo, Ms Huan Yang, Dr. Hongxia Lv and Ms Liying Du for their technical help. We acknowledge the High-performance Computing Platform of Peking University for enabling us to perform data analysis. We acknowledge Dr. Xing Chen and Qi Tang (Peking University) for providing materials and guidance for the experiment of sialic acid detection. We acknowledge Dr. Peng Jiang, Dr. Jie Cheng and Jinxing Yan (Tsinghua University) for providing technical help in mouse experiment. We also acknowledge Dr. Zexian Zeng (Peking University) and Dr. Liang Weng (Xiangya Hospital Central South University) for their valuable advice on this study. This project was supported by funds from the National Natural Science Foundation of China (NSFC 31930016 to W.W.), Beijing Advanced Innovation Center for Genomics at Peking University and the Peking-Tsinghua Center for Life Sciences (to W.W.), and China Postdoctoral Science Foundation (2020M670031, to Y.L.; 2020M670053, to Y.-S.L.).

## AUTHOR CONTRIBUTIONS

W.W. conceived and supervised this project. W.W., Y.L., Y.-S.L., and X.N. designed the experiments. Y.L., Y.-S.L., X.N. and A.C. performed the library screens, all the following validations and experimental data analysis. Y.Y. performed the NGS library construction. Y.-Z.L. performed the bioinformatics analysis. Z.L. provided help for data analysis. Y.L., Y.-S.L., and X.N. wrote the manuscript with the help of A.C. and Y.-Z.L., and W.W. revised it.

## RESOURCE AVAILABILITY

### Lead contact

Further information and requests for resources and reagents should be directed to and will be fulfilled by the Lead Contact, Wensheng Wei.

### Materials availability

All reagents generated in this study are available from the Lead Contact without restriction.

### Data and code availability

The sequence data have been deposited in the Genome Sequence Archive (Chen et al., 2021b) in National Genomics Data Center (Members and Partners, 2022), China National Center for Bioinformation/Beijing Institute of Genomics, Chinese Academy of Sciences (GSA-Human: HRA005746) that are publicly accessible at https://ngdc.cncb.ac.cn/gsa-human. All data supporting the findings in this manuscript are available upon reasonable request.

## EXPERIMENTAL MODEL AND SUBJECT DETAILS

### Mice

The female BALB/c mice and BALB/C nude mice (6 to 8 week old) were ordered from Beijing Vital River Laboratory Animal Technology Co., Ltd. All mice were bred and kept under specific pathogen-free (SPF) conditions in the Laboratory Animal Center of Peking University. The animal experiments were approved by Peking University Laboratory Animal Center (Beijing) and conducted in accordance with the National Institute of Health Guide for Care and Use of Laboratory Animals.

### Cell lines

The HEK293T cell line was obtained from EdiGene Inc., and the A375 and B16F10 cell lines were purchased from ATCC. The A375-ABEmax and B16F10-ABEmax cell lines were generated in this study. HEK293T, A375 and A375-ABEmax cells were cultured in Dulbecco’s modified Eagle’s medium (DMEM, Gibco, #C11965500BT) containing 10% fetal bovine serum (FBS, Biological Industries, #04-010-1A) and 1% penicillin/streptomycin (P/S). B16F10 and B16F10-ABEmax cells were cultured in RPMI1640 medium (Gibco, #C11875500BT) supplemented with 10% FBS and 1% P/S. All cells were cultured with 5% CO_2_ at 37°C and were routinely checked to confirm the absence of mycoplasma contamination using Mycoplasma Detection Kit (InvivoGen, #rep-mys-50).

### Primary human T cells

Peripheral blood mononuclear cells (PBMCs) were obtained from healthy donors with informed consent. Primary human T cells expressing the anti-NY-ESO-1 TCR were generated by retroviral transduction according to previous studies described (Dersh et al., 2021; Patel et al., 2017), and were frozen in the cryopreservation medium (Stemcell Technologies, #100-1061). Once thawed, T cells were maintained in T cell expansion medium (Stemcell Technologies, #10981) supplemented with 10% FBS, 1% penicillin/streptomycin, and 50 ng/mL IL-2 (Stemcell Technologies, #78036.3). T cells were activated and expanded using human CD3/CD28 T cell activator (Stemcell Technologies, #10971) for 3 days, and then subjected to subsequent experiments.

## METHOD DETAILS

### Plasmids

pLenti-ABEmax-P2A-EGFP expression plasmid was constructed by cloning ABEmax_P2A_EGFP sequence from pCMV_ABEmax_P2A_GFP (Addgene, #112101) into the lentiviral vector. All sgRNAs used for validation (Supplementary Table 7) were cloned into the pLenti-sgRNA(lib)-puro vector (Addgene, #119976) through Golden Gate assembly. Protein-coding sequences for cDNA over-expression or co-immunoprecipitation were cloned into pLenti_CMV_cDNA_Flag_SV40_mCherry vector or pLenti_CMV_cDNA_HA_SV40_EGFP vector by PCR and Gibson assembly (NEB, #E2611L). All plasmids were verified by Sanger sequencing.

### ABE screens for functional S/T/Y residues in A375 cells

The A375-ABEmax cells were seeded in 15-cm dishes 24 hours before lentivirus infection, then were respectively transduced with each of the S/T/Y lentiviral libraries (sense library and antisense library) at an MOI of 3 with a high coverage for each sgRNA (about 1,500-fold, about 500-fold for each iBAR). Forty-eight hours post transduction, the library cells were cultured with 1 μg/mL puromycin (Solaribio, #P8230) for two days. After puromycin selection, the time point was denoted as Day 0 of the screening, and the library cells with at least 1,500-fold coverage for sgRNAs were maintained and passaged every 3 days. At Day 10 (IFNγ-absent screens) or Day 13 (IFNγ-treated screens), PD-L1^high/low^ and HLA^high/low^ cells were respectively subjected to the first round of FACS enrichment by BD FACS Aria III or MoFlo Astrios EQ (Beckman). For the PD-L1 screens, cells were pre-treated with or without 100 ng/mL IFNγ (Sino Biological, #GMP-11725-HNAS) for 48 hours and stained with APC anti-human CD274 antibody (BioLegend, #329708) before FACS. For the HLA screens, cells were stained with APC anti-human HLA-A,B,C antibody (BioLegend, #311410) before FACS. At least three times of the library cells were subjected to immunofluorescence staining, and 1 μL antibody per million cells in 100 μL staining buffer (BioLegend, #420201) were used in the staining according to the standard protocol. In each group, the highest and lowest 10% of cells were collected based on APC fluorescence. One week after the first-round sorting, the cells were stained with the same antibodies and were further subjected to FACS enrichment. In the second-round sorting, APC-positive or APC-negative library cells were collected for each group through comparing with A375 cells infected with *AAVS1*-targeted conrol sgRNA (Figure S1B-E). At Day 24, the library cells without FACS were havested as the reference group and the FACS-enriched cells from the second-round sorting of each group were havested as the experimental groups.

### Genomic DNA isolation and amplicon sequencing of the S/T/Y library

Genomic DNA was extracted from reference cells and experimental cells using the DNeasy Blood & Tissue Kit (Qiagen, #69506). For each group, all extracted genomes were used as the PCR templates and the sgRNA-coding sequences with iBAR were amplified using KAPA HiFi HotStart ReadyMix PCR kit (Roche, #KK2631). The DNA amplification was performed under the following condition: 30 s at 95℃ for initial denaturation; 26 cycles consisting of 10 s at 95℃ for denaturation, 30 s at 60℃ for annealing, and 15 s at 72℃ for extension; and 15 s at 72℃ for final extension. The PCR products of each group were pooled and purified using DNA Clean & Concentrator-25 kit (Zymo Research, #D4034), followed by next-generation sequencing (NGS) analysis on Illumina HiSeq X TEN platform.

### Computational analysis of screens

To analyze the NGS data of the screens, we used MAGeCK-iBAR algorithm (Zhu et al., 2019) to evaluate the change of sgRNA abundance between the reference group and each experimental group. We used default parameters of MAGeCK-iBAR to calculate the *p* value (lo_value in the output) for each sgRNA considering both the significance and consistency of three iBARs. The final screen score was defined as -log10 of *p* value after Benjamini-Hochberg (BH) adjustment and sgRNAs with a screen score of more than 1 were selected as the negatively or positively enriched candidates for follow-up studies.

### Validation of candidate sites identified from the screens

A375-ABEmax cells were transduced with lentivirus of each sgRNA targeting candidate site or *AAVS1* at an MOI of >1, and the time point of lentivirus infection was denoted as T0. Forty-eight hours post transduction (T2), cells were treated with 1 μg/mL puromycin for two days and the resistant cells were passaged for two generations. For the IFNγ-absent condition, sgRNA-infected cells were collected at T9. For the IFNγ-treated condition, sgRNA-infected cells were seeded at T8, treated with 100 ng/mL IFNγ at T9 for 48 h, and finally collected at T11. For both conditions, sgRNA-infected cells cultured in 6-well plates were washed by DPBS (Gibco, #C14190500BT), followed by detachment using accutase (BioLegend, #423201). One million cells were collected and resuspended in 100 μL staining buffer with 1 μL APC anti-human CD274 antibody or APC anti-human HLA-A,B,C antibody following the standard protocol. Flow cytometry analysis was performed with the BD LSRFortessa SORP (BD Biosciences). Changes in PD-L1 or MHC-I surface expression were calculated as the changes in raw median fluorescence intensity (MFI). The relative MFI of all samples was normalized to the isotype control or further normalized to the *AAVS1*-targeted control cells. Antibodies used in the validation include anti-human CD274 antibody (APC, clone 29E.2A3, BioLegend, #329708), anti-human HLA-A,B,C antibody (APC, clone W6/32, BioLegend, #311410), anti-human HLA-BC antibody (APC, clone B1.23.2, eBioscience™, #17-5935-42), Mouse IgG2b, κ Isotype Ctrl (APC, clone MPC-11, BioLegend, #982108), Mouse IgG 2a, κ Isotype Ctrl (APC, clone MOPC-173.

### Detection of base editing outcomes by NGS

A375-ABEmax or B16F10-ABEmax cells were transduced with lentivirus of each sgRNA targeting candidate site at an MOI of >1, and were further treated with puromycin as described above. Seven days post transduction, sgRNA-infected cells were collected and subjected to genome DNA isolation. For the mutant cells and WT cells, about 200-bp genomic sequences surrounding each sgRNA-targeted site were amplified using specific primers by PrimeSTAR® GXL Premix (TAKARA, #R051A), followed by NGS analysis on Illumina HiSeq X TEN platform. The paired-end NGS data was first assembled by PANDAseq software. The sequence of sgRNA-targeted regions was extracted from the assembled fasta files by their flanking sequence, which was 10 bp upstream and 10 bp downstream of the sgRNA-targeted regions. The percentage of A/T/C/G in each position was further calculated, including the targeted site and sgRNA editing window, to assess on-target editing efficiency as well as bystander editing for each candidate sgRNA.

### Real-time qPCR analysis

For the IFNγ-absent or IFNγ-treated condition, A375-ABEmax cells infected with each indicated sgRNA were respectively collected at T9 or T11 as described above. RNA of the sgRNA-infected cells was extracted using RNAprep pure Cell/Bacteria Kit (TIANGEN, #DP430), and the cDNA was synthesized using HifairII 1st Strand cDNA Synthesis SuperMix (YEASEN, #11120ES60). Real-time qPCR was performed using TB Green Premix Ex Taq II (TaKaRa, #RR820A) on Roche LightCycler480 Real-Time PCR System. All cDNA samples were assayed in triplicate and the relative RNA expression level of each sample was normalized by *GAPDH*. All the primers used for real-time qPCR are listed in Supplementary Table 8.

### Immunoblotting

A375-ABEmax cells infected with each indicated sgRNA were inoculated in 6-well plates, and were respectively collected at T9 or T11 for different IFNγ treatments as described above. Cells were washed twice with PBS, and were lysed using pre-cooled RIPA lysis buffer supplemented with protease and phosphatase inhibitor (Thermo Fisher Scientific, #78441) on ice for 30 min. After quantifying the protein concentration by the BCA method (Thermo Fisher Scientific, #23225), the lysates were electrophoretically separated by 12% SDS-PAGE gel and transferred to a PVDF membrane (Bio-Rad, #10026934). The proteins were blocked with 5% skim milk (Thermo Fisher Scientific, #232100) in PBST or TBST at room temperature for 1 h and were further incubated with the primary antibody at 4 ℃ overnight. The PVDF membranes were washed with PBST or TBST three times and then incubated with HRP secondary antibodies (1:10000) at room temperature for 1 h. The secondary antibodies includes: goat anti-rabbit IgG-HRP (Jackson Immunoresearch, #111035003) or goat anti-mouse IgG-HRP secondary antibody (Jackson Immunoresearch, #115035003). After being washed with TBST three times, the protein bands were detected by using Clarity^TM^ Western ECL Substrate Kit (Bio-Rad, #1705060) on the Chemidoc^TM^ system (Bio-Rad, #1708370).

### Immunoprecipitation

8 × 10^5^ HEK293T cells were seeded in 6-well plates for each sample. The cells were transfected with indicated plasmids on the second day, followed by stimulation with 100 ng/mL IFNγ on the third day for 48 h. Cells were washed in PBS, lysed in RIPA lysis buffer with protease and phosphatase inhibitor on ice for 30 min, and further pelleted by centrifugation at 12,000 g for at 4 °C for 10 min. The supernatant was collected with 30 µL of the cell lysates as the input, and the rest was treated with Anti-Flag M2 Affinity Gel (Sigma-Aldrich, #A2220) or HA beads (Sigma-Aldrich, #E6779) at 4°C overnight. After washing the lysates four times with RIPA buffer, 5 × loading buffer was added to the sample, followed by boiling at 100 °C for 10 min. Then the immunoblotting analysis was carried out as described above. Antibodies used for immunoblotting include: rabbit polyclonal anti-HA (Sigma-Aldrich, #H6908/SAB4300603, 1:10000), rabbit polyclonal anti-FLAG (Sigma-Aldrich, #F7425, 1:10000) and mouse monoclonal anti-FLAG (Sigma-Aldrich, #F1804, 1:10000).

### Detection of sialic acid by flow cytometry

A375-ABEmax cells infected with sgRNA targeting SLC35A1_Y98 and *AAVS1* were collected at the 9th day post lentivirus infection. After DPBS washing and accutase detachment, one million cells were washed by DPBS twice, resuspended in 1 mL PBS supplemented with 0.5% BSA. The Maackia Amurensis Lectin II (MAL-II)-biotin (Vector Laboratories, #B-1265) was added to the suspension at a final concentration of 5 µg/mL, followed by incubation at room temperature for 30 min. Next, the cells were washed with DPBS three times and stained with 1 µg/mL Streptavidin-Alexa Fluor 647 (AF647) (BioLegend, #405237) for another 30 min. After washing with DPBS three times, flow cytometry analysis was performed to detect the AF647 (APC) signal with the BD LSRFortessa SORP (BD Biosciences). Changes in sialic acid surface expression were calculated as the changes in raw MFI, and the relative MFI was generted by normalization to the fluresence of unstained cells.

### Competative T cell killing assay

A375 cells, which endogenously express NY-ESO-1 antigen, were further engineered to stably overexpress ABEmax with an EGFP marker in this study. In the co-culture experiment, A375-ABEmax cells infected with each indicated sgRNA were first mixed with A375 WT cells in a 1:1 ratio, then were seeded in 48-well plates and allowed to attach for 12 h before adding the anti-NY-ESO-1 TCR-transduced primary human T cells at an appropriate effector to target cell (E:T) ratio. Meanwhile, paired controls without adding T cells were included for each condition. After co-culturing the targeted A375 cells and T cells in RPMI 1640 medium (Gibco, #11875093) for 6 h, the cells were washed twice by DPBS to remove most of the surface T cells. Then the A375 cells along with some adherent T cells were detached with accutase, followed by staining with anti-human CD3 (UCHT1, BV650, BioLegend, #00467) and DAPI (BioLegend, #422801) to further exclude T cells and dead cells. Flow cytometry analysis was performed with the BD LSRFortessa SORP (BD Biosciences), and the percentage of EGFP+ cells was measured after gating out T cells and dead cells. The extent of the killing sensitivity was defined as: 100 ×[1-(A1/100-A1)/(B1/100-B1)], A1: Percentage of A375-ABEmax cells (represented as EGFP+ cells) that were incubated with T cells, B1: Percentage of A375-ABEmax cells that were not incubated with T cells. The extent of the killing resistance was defined as: 100×[1-(A2/100-A2)/(B2/100-B2)], A2: Percentage of A375 WT cells that were incubated with T cells, B2: Percentage of A375 WT cells that were not incubated with T cells (Joncker et al., 2010). For each sample, both of the co-culture assay and the paired control were performed in triplicate.

### RNA-seq and data analysis

The sgRNA targeting NAA20_Y137, SETD2_Y1666 or *AAVS1* was individually transduced into A375-ABEmax cells at an MOI of <1 in duplicate or triplicate. At T11 as described above, 2 × 10^6^ cells were collected after IFNγ treatment for two days. The total RNA of each sample was extracted using the RNeasy Mini Kit (QIAGEN, #79254), and the RNA-seq libraries were prepared as previously described (Ding et al., 2021). All samples were subjected to NGS analysis using the Illumina HiSeq X TEN platform. The RNA sequencing data was first processed by FASTP software to cut adapters and filter low quality sequences. Then HISAT2 was used to map the reads to human reference genome hg38 under default parameters. The raw counts of mapped reads for each gene were calculated using featurecounts software. The annotation file for this step was from GENCODE v38 gtf file and the reads in exon level (-t parameter) were counted. The differential gene expression analysis was performed by DESeq2 package (V1.40.2) and the downstream GO enrichment was performed by clusterProfiler package (V3.10.1).

### Chromatin immunoprecipitation with sequencing (ChIP-seq) and data analysis

The ChIP assays were performed using Hyperactive Universal CUT&Tag Assay Kit for Illumina (Vazyme, #TD903). The procedure was according to manufacturer’s instructions. Breifly, sgRNA targeting SETD2_Y1666 or *AAVS1* was individually transduced into A375-ABEmax cells at an MOI of <1 in triplicate, and 50,000 cells were harvested at T11 after IFNγ treatment for two days. Cells were fixed on cleaned NovoNGS CoA beads, followed by incubation with primary anti-H3K36me3 antibody (Abcam, #ab9050) at 4 ℃ overnight. On the next day, Immunoprecipitates was incubated with Goat anti-Rabbit IgG antibody (1:100) at room temperature for 30 min, and further incubated with protein A/G-Tn5 transposase and ChiTag buffer for 1 h. Next, the samples were subjected to DNA fragementation by adding tagmentation buffer with incubation at 37 ℃ for 1 h, followed by DNA extraction through incubation with tagment DNA extract beads, thus obtaining fragmented DNA. Then the ChIP samples were prepared for NGS analysis using VAHTS Universal DNA Library Prep Kit for Illumina v.3 (Vazyme, #ND607) and deep-sequenced on the Illumina HiSeq X TEN platform. The cleaned fastq files was first mapped to human reference genome hg38 using BOWTIE2 under default parameters. Then we used MASC2 to call peaks and chose broad peak pattern considering features of H3K36me3. Different peak analysis was performed by DiffBind package (V2.10.0) in R. Integrative Genomics Viewer (IGV) was used to visualize peaks in the interested regions and the results from three replicates were merged in IGV.

### Mouse experiments

For the immune-competent mouse model, sgRNA targeting Naa20_Y137, Setd2_Y1640 and the negative control sgRNA was individually transduced into B16F10-ABEmax cells, then 4×10^5^ sgRNA-infected cells were subcutaneously inoculated into the right flank of 6-8 week-old female C57BL/6 mice, which were further divided into control or experimental groups randomly. From Day 7 post transplantation when the tumor volume reached about 100 mm^3^, the control and experimental groups were treated with control IgG (BioXcell, #BE0089, 200 μg per mouse) or anti-PD-1 (BioXcell, #BE0273, 200 μg per mouse) by intraperitoneal injection every three days for a total of four times (on the 7th, 10th, 13th and 16th days), and monitor of the tumor growth was finished on the 19th day. For the immunodeficient mouse model, 2×10^5^ B16F10 cells infected with each indicated sgRNA were subcutaneously inoculated into the right flank of 6-week-old female BALB/c nude mice. Tumor growth was measured using digital calipers, and tumor sizes were recorded every three days until the sizes reached 2000 mm^3^.

### Isolation of the tumor infiltrated immune cells and flow cytometry analysis

The mouse tumor samples separated from the mice were washed with PBS, then were minced into small pieces and further digested by the RPMI 1640 medium supplemented with 1 mg/mL collagenase D (OKA, #D10032) at 37 °C for 30 min. After terminating the digestion by adding RPMI 1640 medium supplemented with 10% FBS, the solutions were filtered through a 200-mesh cell sieve and centrifuged at 260 g for 4 min. Then the cell pellets were washed by PBS and centrifuged at 260 g for 4 min, thus obtaining single-cell suspensions. Cells were stimulated with anti-CD3/CD28 (3.5 µg/ml anti-CD3 mAb, BioLegend, #100339; 1 µg/ml anti-CD28 mAb, BioLegend, #102115) in the presence of 5 μg/mL Brefeldin A (BFA, Thermo Fisher Scientific, #00-4506-51) and 5 μg/mL monensin (Thermo Fisher Scientific, #00-4505-51), and cultured in a humidified incubator with 95% air/5% CO_2_ at 37°C incubation for 4 hours. Cells were collected by centrifugation at 260 g for 4 min, then washed with 1 mL PBS. After centrifugation at 260 g for 4 min to remove the supernatant, the cells were first stained with anti-CD8a mAb (PE, clone 53-6.7, BioLegend, #100708), then fixed with 2× IC fixation buffer (Thermo Fisher Scientific, #00-8222-49) at room temperature for 15 min in the dark, and treated with 1× permeabilization buffer (Thermo Fisher Scientific, #00-8333-56). After centrifugation at 5,000 g for 2 min, the cell pellets were stained with anti-GzmB antibody (FITC, clone QA16A02, BioLegend, #372206), followed by flow cytometry analysis.

## QUANTIFICATION AND STATISTICAL ANALYSIS

### Generation of the SETD2_Y1666-mutation signature

The SETD2-Y1666 mutation signature was defined by extracting top 250 upregulated and top 250 downregulated genes and using the normalized DESeq2 wald statstics as weights, which were calculated on the basis of the equation *k*_*i*_ = *w*_*i*_/*max*(*w*). The *k_i_* stands for the weight of the *i th* gene and *w_i_* indicated the wald statistics of the *i th* gene. Each input expression profiles then could be assessed by computing a SETD2-Y1666 mutation signature score by calculating the sum expression level of the signature genes following the equation *S* = ∑*n*_*i*_ = (*k*_*i*_ ∗ *X*_*i*_), where *S* denotes the signature score and *X*_*i*_ denotes the expression level of the *i t*h gene.

### Immunotherapy trials used for correlation analysis

We collected 91 RNA-seq expression profiles from 54 melanoma patients who were treated with anti-PD-1 therapy from published study (Gide et al., 2019). For each RNA-seq sample, the gene expression profile was analyzed following standard pipeline as described above.

### Correlation analysis between the SETD2_Y1666-mutation signature and representative markers

Referred to the melanoma patients’ cohort that we used (Gide et al., 2019), the MHC-I expression levels were calculated as the average log_2_TPM of *HLA-A*, *HLA-B*, *HLA-C*, and *B2M*, the PD-L1 expression levels were calculated as the average log_2_TPM of *CD274*, and the CTL (cytotoxic T lymphocyte) expression levels were calculated as the average log_2_TPM of *CD8A*, *CD8B*, *GzmA*, *GzmB*, and *PRF1*. The Pearson correlations were computed between the SETD2_Y1666-mutation signature and the expression levels of MHC-I, PD-L1, and CTL.

### Survival analysis

The clinical relevance of SETD2_Y1666 in regulating ICB response was confirmed by testing the association between SETD2_Y1666-mutation signature and progressive survival of patients in immunotherapy trials with cox regression.

### Statistical analysis

Statistical tests, exact value and description of n were presented as described in the figure legends. Unless otherwise noted, n represents biological replicates of the samples (e.g., independent cell cultures, individual tumors, *etc*.). The statistical significance was evaluated using Student’s *t* test or two-way ANOVA (with BH adjustment for multiple testing), and determined as *P* < 0.05, labeled as \**P* < 0.05, \*\**P* < 0.01, \*\*\**P* < 0.001, \*\*\*\**P* < 0.0001.

